# Nutrigenomic Regulation of Sensory Plasticity

**DOI:** 10.1101/2021.12.17.473205

**Authors:** Hayeon Sung, Anoumid Vaziri, Daniel Wilinski, Riley K.R. Woerner, Peter L. Freddolino, Monica Dus

## Abstract

Diet profoundly influences brain physiology, but how nutritional information is transmuted into neural activity and behavior changes remains elusive. Here we show that the metabolic enzyme O-GlcNAc Transferase (OGT) moonlights on the chromatin of the *D. melanogaster* gustatory neurons to instruct changes in chromatin accessibility and transcription that underlie sensory adaptations to a high sugar diet. OGT works synergistically with the Mitogen Activated Kinase/Extracellular signal Regulated Kinase (MAPK/ERK) *rolled* and its effector *stripe* (also known as EGR2 or Krox20) to integrate activity information. OGT also cooperates with the epigenetic silencer *Polycomb Repressive Complex 2.1* (PRC2.1) to decrease chromatin accessibility and repress transcription in the high sugar diet. This integration of nutritional and activity information changes the taste neurons’ responses to sugar and flies’ ability to sense sweetness. Our findings reveal how nutrigenomic signaling generates cell-specific responses to global nutrient variations.

## INTRODUCTION

The levels and types of dietary nutrients play an essential role in cellular processes such as growth, division, and differentiation by providing fuel and biomass. However, nutrients can also affect these aspects of cell physiology by influencing, and often orchestrating, gene expression programs (Dai et al., 2020; Vaziri and Dus, 2021). These effects are mediated through nutrient-sensitive modifications to DNA, RNA, and proteins, as well as changes to the activity, binding, and localization of enzymes and signaling factors (Huang et al., 2015; Katada et al., 2012). These nutrigenomic signaling pathways –nutrigenomics is the field that studies food-genes interactions– could explain how the food environment affects the risk of non-communicable diseases such as diabetes, cancer, and neurodegeneration. They also hold the potential to uncover new interventions and treatments for these debilitating diseases. While the effects of nutrients on gene expression are well-established, relatively little is known about the molecular mechanisms at the food-gene interface. A significant challenge of the field has been to explain how global variations in the nutrient environment lead to specific changes in cell physiology. To overcome these challenges, we have developed an experimental system where the contributions of nutrients to physiology can be studied mechanistically and *in vivo (Vaziri and Dus, 2021)*. Here we use this model to characterize how cell-specific signals are integrated with broad changes in metabolism to drive adaptations to the dietary environment.

Taste sensation changes depending on diet composition. In animals, the levels of bitter, sweet, and salty foods influence how these taste stimuli are perceived, with a general inverse relationship between the amount of a particular food in the diet and the responses of the sensory system to it (May and Dus, 2021; Reed et al., 2020; Sarangi and Dus, 2021). For example, in humans and rodents, the dietary concentration of sugars affects sweetness intensity or the electrophysiological responses of the sensory nerves to sucrose (May and Dus, 2021; McCluskey et al., 2020; Sartor et al., 2011; Sung et al., 2022; Wise et al., 2016). A similar phenomenon occurs in flies, where diets supplemented with 15-30% sucrose, glucose, or fructose decrease the responses of the sensory neurons to sucrose and the transmission of the sweetness signal to higher brain areas <u>(May et al. 2019; Vaziri et al. 2020; Wang et al. 2020; Ganguly et al. 2021;</u> <u>May et al. 2020)</u>. In rats and flies, the dulling of the sensory system to sugar occurs even without weight gain, suggesting that diet exposure is sufficient to drive sweet taste plasticity <u>(Sung et al.</u> <u>2022; May et al. 2019)</u>. Our previous work in flies implicated metabolic signaling through the Hexosamine Biosynthesis Pathway (HBP) enzyme O-GlcNaC Transferase (OGT) in this phenomenon (May et al., 2019). Specifically, knockdown of OGT exclusively in the fly sweet-taste cells prevented the neural and behavioral decrease in sugar responses observed with a high-sugar diet (May et al., 2019). OGT uses the metabolic end-product of the HBP, UDP-GlcNAc, to post-translationally modify proteins and change their stability or activity (Hart, 2019). OGT activity is sensitive to all cellular levels of UDP-GlcNAc without substrate inhibition, but it is enhanced by high dietary sugar due to a higher flux through the HBP (Bouché et al., 2004; Hanover et al., 2010; Hawkins et al., 1997; Marshall et al., 2004; May et al., 2019; Na et al., 2015; Olivier-Van Stichelen et al., 2017; Wang et al., 1998; Wilinski et al., 2019). OGT is also a nucleocytoplasmic protein that interacts with many chromatin- and DNA-modifying complexes; as such, it is thought to function as a nutrigenomic sensor, bridging diet and genes (Hardivillé and Hart, 2014; Hart, 2019; Olivier-Van Stichelen et al., 2017; Olivier-Van Stichelen and Hanover, 2015). Despite global changes in HBP flux with high dietary sugar, the consequences of OGT activity differ among cell types. Understanding how this occurs would provide an opportunity to uncover how specificity is achieved in nutrigenomic signaling. Here we exploited the effects of OGT on *Drosophila* sensory neurons and the exquisite genetic tools of this organism to investigate this question. Our experiments reveal that nutrigenomic specificity is achieved through integrating metabolic state with cell-specific experience. In the sensory neurons, OGT decorates nutrient-sensitive loci also occupied by the epigenetic silencer *PRC2.1* and the activity-dependent ERK effector Stripe (*Sr*). This cooperation leads to changes in chromatin accessibility and transcription that drive sensory plasticity, and the catalytic activity of OGT plays an instructional role in this process. Thus, our results uncover mechanistic insights into how nutrigenomic signaling translates nutritional information into cell-specific adaptations.

## RESULTS

### The nutrient sensor OGT decorates the chromatin of sweet sensory cells

Since transcriptional changes have been implicated in sugar diet-induced taste plasticity (May et al., 2019; Vaziri et al., 2020; Wang et al., 2020) and OGT associates with chromatin binding factors (Gambetta and Müller, 2015; Gao et al., 2018; Hart et al., 2011; Vella et al., 2013), we asked whether this metabolic enzyme moonlights on the chromatin of sweet taste neurons. We used <u>D</u>NA <u>a</u>denosine <u>m</u>ethyltransferase <u>Id</u>entification (Dam-ID or TaDA) to measure the association of OGT with DNA (Marshall et al., 2016; van Steensel and Henikoff, 2000) and <u>C</u>hromatin <u>A</u>ccessibility profiling using <u>Ta</u>rgeted <u>Da</u>mID (CaTaDA) to assess chromatin accessibility (Sen et al., 2019). Transgenic UAS-*LT3-Dam::OGT* or UAS-LT3-*Dam* flies were crossed with *Gustatory Receptor 5a* GAL4 (*Gr5a*) flies (Chyb et al., 2003) to drive expression exclusively in the ~60 sweet taste cells of the fly mouthpart, and *Tubulin-GAL80ts* to control the timing of transgene induction. *Gr5a>LT3-Dam::OGT; tubulin-GAL80ts* (green) and *Gr5a>LT3-Dam; tubulin-GAL80ts* (yellow) transgenic flies were kept at the permissive temperature and fed a control (CD, 5% sucrose) or sugar (SD, 30% sucrose) diet for 3 days. *Dam::OGT* and *Dam* were then induced by heat shocking the animals at 30°C for 18 hours on day 4, as in our prior experimental design (Fig. 1A) (Vaziri et al., 2020). The normalized *Dam::OGT* replicates clustered together by diet (Fig. S1A), and the chromatin accessibility profile of Dam at the *Gr5a* sweet taste receptor gene promoter was high, while at the bitter *Gustatory Receptor Gr66a* promoter– only expressed in adjacent cells– accessibility was low (Fig. S1B), suggesting that these transgenes were targeted to the correct cells.

**Figure 1:**
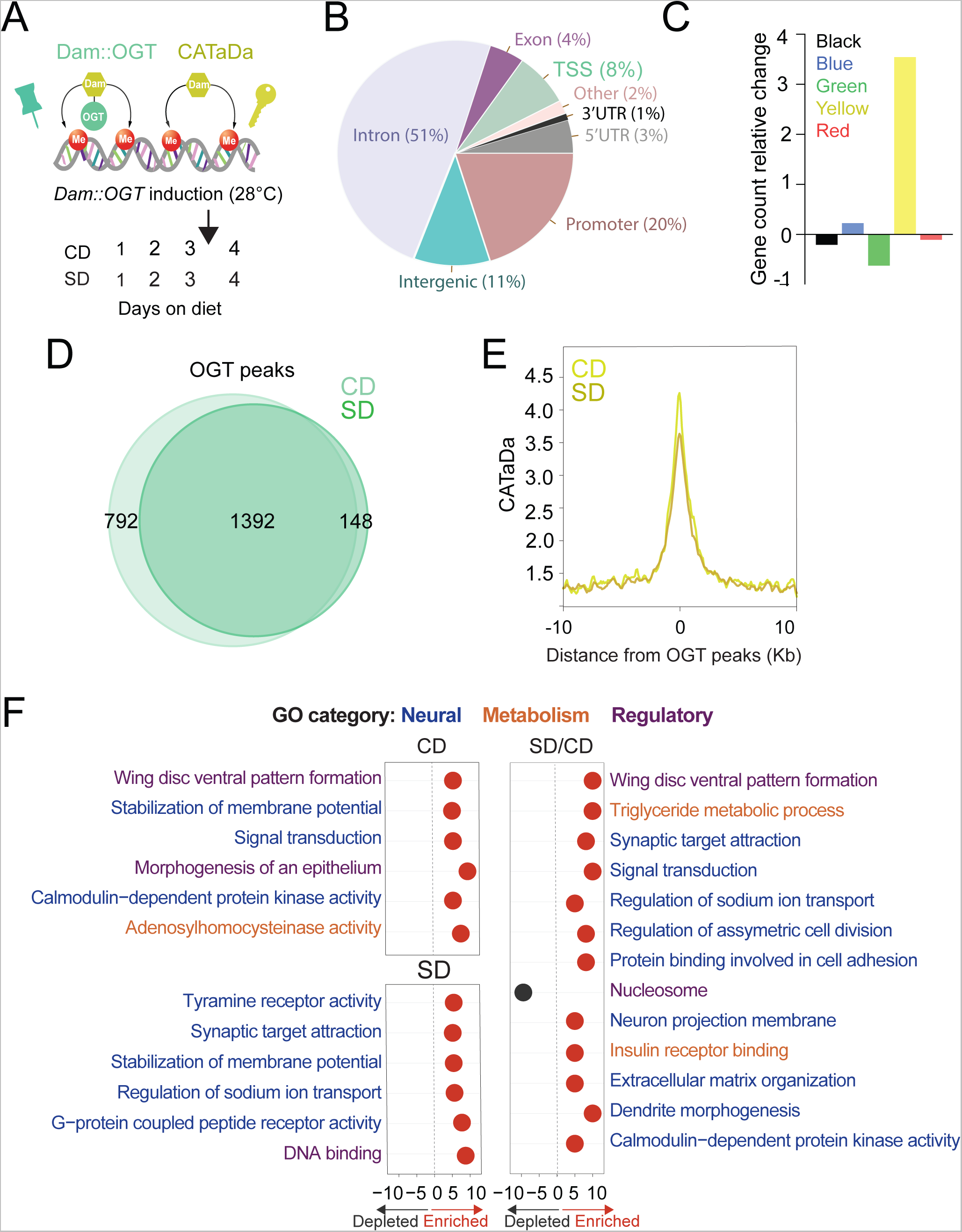
OGT decorates the chromatin of the sweet taste cells. **(A)** Design of Targeted Dam-ID for OGT occupancy (*Dam::OGT*) and *Dam* accessibility (CATaDa) experiments. Age-matched *Gr5a*;*tubulin-GAL80^ts^ > UAS-LT3-Dam::OGT* and *Gr5a*;*tubulin-GAL80^ts^>UAS-LT3-Dam* flies were placed on a CD or SD for 3 days at 20° to 21°C and then switched to 28°C between days 3 and 4 to induce expression of the transgenes. **(B)** Annotation of OGT chromatin occupied regions using HOMER. **(C)** The proportion of observed *Dam::OGT* consensus peaks allocated to their respective chromatin domains normalized to the expected proportions across the whole genome. **(D)** Overlap of log2*(Dam::OGT/Dam)* chromatin binding peaks of CD (light green) and SD (dark green) (find_peaks, q<0.01). **(E)** Average CATaDa signal on a CD (light yellow) and SD (dark yellow) centered at OGT peaks. **(F)** iPAGE summary plots for OGT peaks on a CD (top left), SD (bottom left), and the difference of SD/CD (right). Text in blue represents neural GO terms, orange represents metabolic GO terms, and purple represents regulatory GO terms.

Dam::OGT was associated with chromatin at introns (51%) and Transcriptional Start Sites (TSS) and promoters (30%) (Fig. 1B, Supplemental File 1); these patterns are similar to those observed in the only other study that measured OGT occupancy on the chromatin of mouse embryonic stem cells (Vella et al., 2013). In flies, chromatin types are classified according to the histone modifications present and the proteins bound (Filion et al., 2010). Using this chromatin characterization, we found that OGT was enriched in transcriptionally active euchromatin, characterized by H3K4 and H3K36 methylation (also known as “yellow”) (Fig. 1C). OGT was also enriched at “blue” Polycomb heterochromatin decorated by H3K27 methylation and bound by PRC2 (Fig. 1C) (Filion et al., 2010; Vaziri et al., 2020).

We next examined the differential binding of OGT between the two diets. Although the majority of intervals were shared between a CD and SD (Fig. 1D, find_peaks FDR <0.01), a few hundred loci were uniquely associated with OGT in either the CD (36%) or SD (10%) condition. However, the chromatin accessibility at OGT-bound peaks decreased in the high sugar diet condition (Fig. 1E). To characterize the function of the genes occupied by OGT, we performed pathway enrichment analysis using iPAGE (Goodarzi et al., 2009). On CD, OGT-decorated genes were involved in signal transduction, membrane potential, and calmodulin-dependent protein kinase activity (Fig. 1F, left). Instead, genes targeted by OGT in the SD condition were enriched in G-protein coupled receptor activity, synaptic target attraction, and transcription (Fig. 1F, left). Finally, genes with differential OGT binding were enriched for regulatory/signaling and neural GO terms, including dendrite morphogenesis, neuron projection membrane, synaptic target attraction, signal transduction, pattern formation, and asymmetric cell division (Fig. 1F right, for full iPAGE, GO term analysis see Fig. S2). Together, these experiments show that OGT resides on the chromatin of the sweet taste at open domains characterized by a small, but significant diet sensitivity, and that genes associated with neural functions are particularly abundant among the set with diet-dependent OGT binding.

### OGT and PRC2.1 share diet-sensitive chromatin sites

Our previous work showed that the epigenetic silencer PRC2.1– specifically its histone 3 lysine 27 methyltransferase activity– was necessary and sufficient to drive sweet taste plasticity in response to the nutrient environment (Vaziri et al., 2020). In the presence of high dietary sugar, PRC2.1 decreased chromatin accessibility and expression of transcription factors involved in synaptic function and signaling. Silencing these genes and their regulons lowered neural and behavioral responses to sweetness in high-sugar diet flies (Vaziri et al., 2020). Since OGT and PRC2.1 play a role in sweet taste plasticity and OGT occupancy was enriched at blue Polycomb chromatin, we asked whether there was an overlap in their occupancy.

A comparison of the peaks occupied by both *Dam::Pcl* (*pink*, *Pcl* is the recruiter for PRC2.1) and *Dam::OGT* (green) revealed a small number of shared intervals (Fig. 2A, ~10%; Supplemental File 1). These 162 loci were enriched in the blue “Polycomb” chromatin (p<0.001, permutation test) and had lower expression levels in the *Gr5a+* neurons (from TRAP experiment in (Vaziri et al., 2020)) compared to those bound by OGT alone (Fig. 2B, *purple* vs. *green*) (Fig. 1C) (Filion et al., 2010). OGT x Pcl intervals had higher expression than those occupied by PRC2.1 alone, suggesting they may represent a subtype of Polycomb chromatin (Fig. 2B, *purple* vs. *pink*). We next asked whether the dietary environment changed the association of OGT and Pcl at these loci. There was more OGT and Pcl at the OGT x Pcl shared sites in the SD condition compared to CD, and more OGT than Pcl was present at these sites in both diets (Fig. 2C). Strikingly, chromatin accessibility at OGT x Pcl was markedly (50%) decreased on SD compared to CD (Fig. 2D). This nutrient-dependent shift in accessibility was 3-fold higher at the shared loci compared to those bound by OGT alone (compare Figs. 1E and 2D; also comparatively higher than those bound by PRC2.1 alone (Vaziri et al., 2020)).

**Figure 2:**
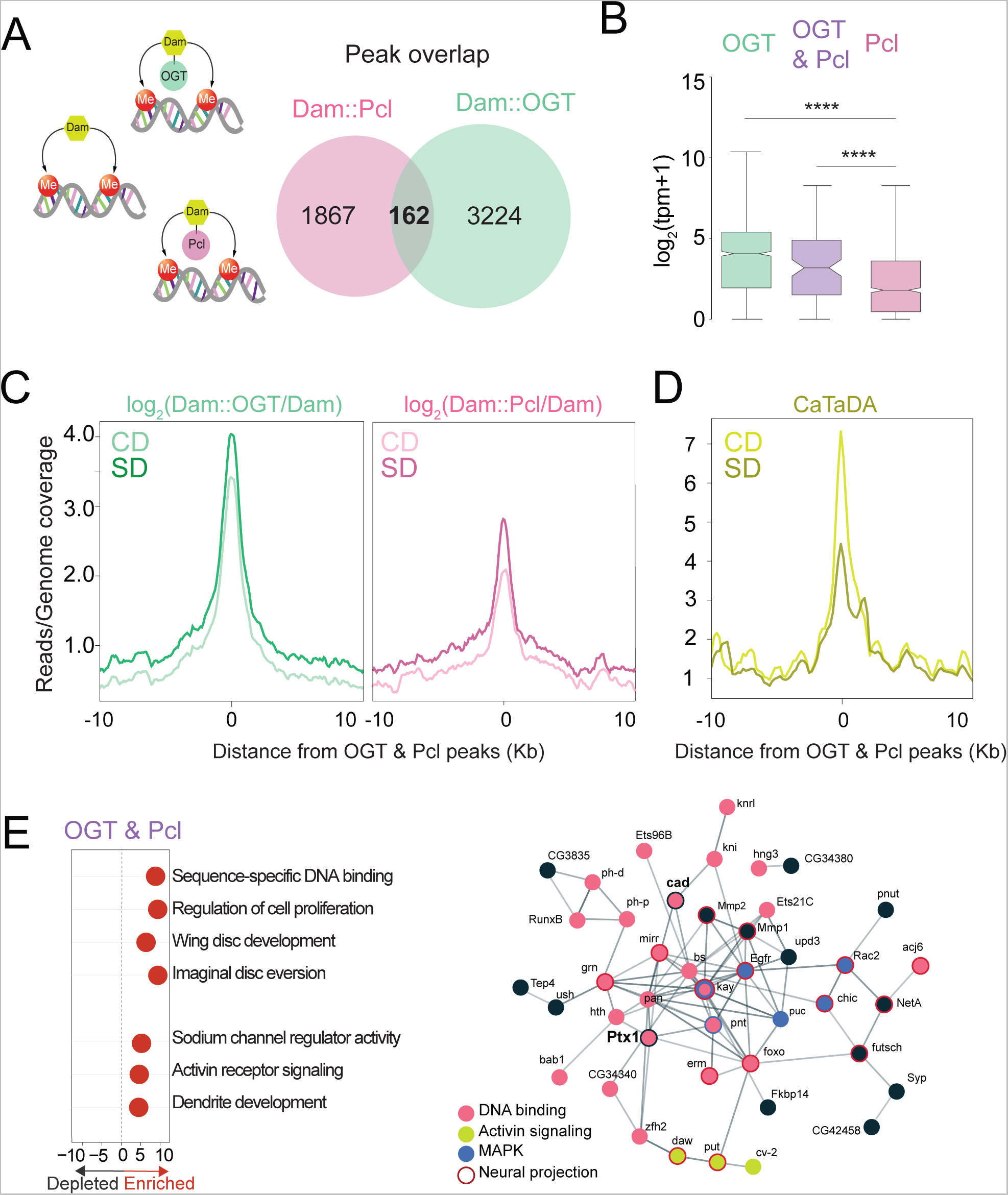
OGT and PRC2.1 mark nutrient-sensitive chromatin in the sweet taste cells. **(A)** Diagram of the Targeted Dam-ID (TaDa occupancy, *Dam::OGT* green, and *Dam::Pcl* pink) and (CATaDa, accessibility, yellow) experiments analyzed in this figure. Overlap of log2(*Dam::Pcl/Dam*, pink) and log2(*Dam::OGT/Dam*, green) chromatin occupancy peaks (peak calling: find_peaks, q<0.01). **(B)** The distribution of the TPMs of genes occupied by OGT (green), Pcl (pink), and genes co-occupied by OGT and Pcl (purple). **(C)** Average log2(*Dam::OGT/Dam*; left) and log2(*Dam::Pcl/Dam*) (right) signal on a CD (lighter shades) and SD (darker shades) diet centered at OGT + Pcl co-occupied peaks. **(D)** Average CATaDa signal on a CD (lighter shade) and SD (darker shade) centered at OGT + Pcl co-occupied peaks. **(E) (left)** iPAGE pathway analysis of genes co-occupied by OGT and Pcl and (right) STRING interaction network of genes co-occupied by OGT+Pcl, colors represent GO terms from the pathway enrichment analysis.

GO term analysis of genes shared by OGT/Pcl targets revealed enrichment in regulatory pathways involved in sequence-specific DNA binding, including those implicated in neural differentiation, sodium channel regulator activity, Activin receptor signaling, and dendrite development (Fig. 2E, left). 30% of the OGT x Pcl sites corresponded to genes encoding DNA-binding and regulatory factors, including two Homeobox transcription factors known to play a role in sweet taste function and plasticity, *cad* and *Ptx1* (Fig. 2E, right) (Vaziri et al., 2020). Analysis of protein interactions between OGT x PRC2.1 genes (Szklarczyk et al., 2020) uncovered a Protein-Protein Interaction network enrichment (*p<1.0e-16*) among DNA-binding factors (pink, *p=2.08e-09*), Mitogen-Activated Protein Kinase (MAPK, blue, *p=0.00059*), signal transduction (Tumor Growth Factor TGF-β/Activin signaling, yellow), neuron projection (red outline, *p=4.95e-7*), and response to stimuli (p=7.15e-0.5). Interestingly, the MAPK pathway is sensitive to neural activity and nutrients (Papa et al., 2019; Robles-Flores et al., 2021).

### The catalytic activity of OGT is required for all aspects of diet-induced taste plasticity

Our data show that OGT moonlights on the chromatin of the sensory neurons and that its binding is diet-dependent at loci also occupied by PRC2.1. To understand the effects of OGT function, and thus nutrigenomic signaling, on taste plasticity, we inhibited the catalytic activity of OGT using the OGT Small Molecule Inhibitor-1 (OSMI) (May et al., 2020; Ortiz-Meoz et al., 2015)(Fig. 3A). Although OSMI treatment is not specific to sweet taste cells, this approach has several advantages: 1) it only affects the catalytic activity of OGT, not its protein levels; 2) it allows us to manipulate OGT activity only during the dietary exposure period, and 3) it makes experiments that would be otherwise inaccessible because of complex genetic combinations and lethality possible.

**Figure 3:**
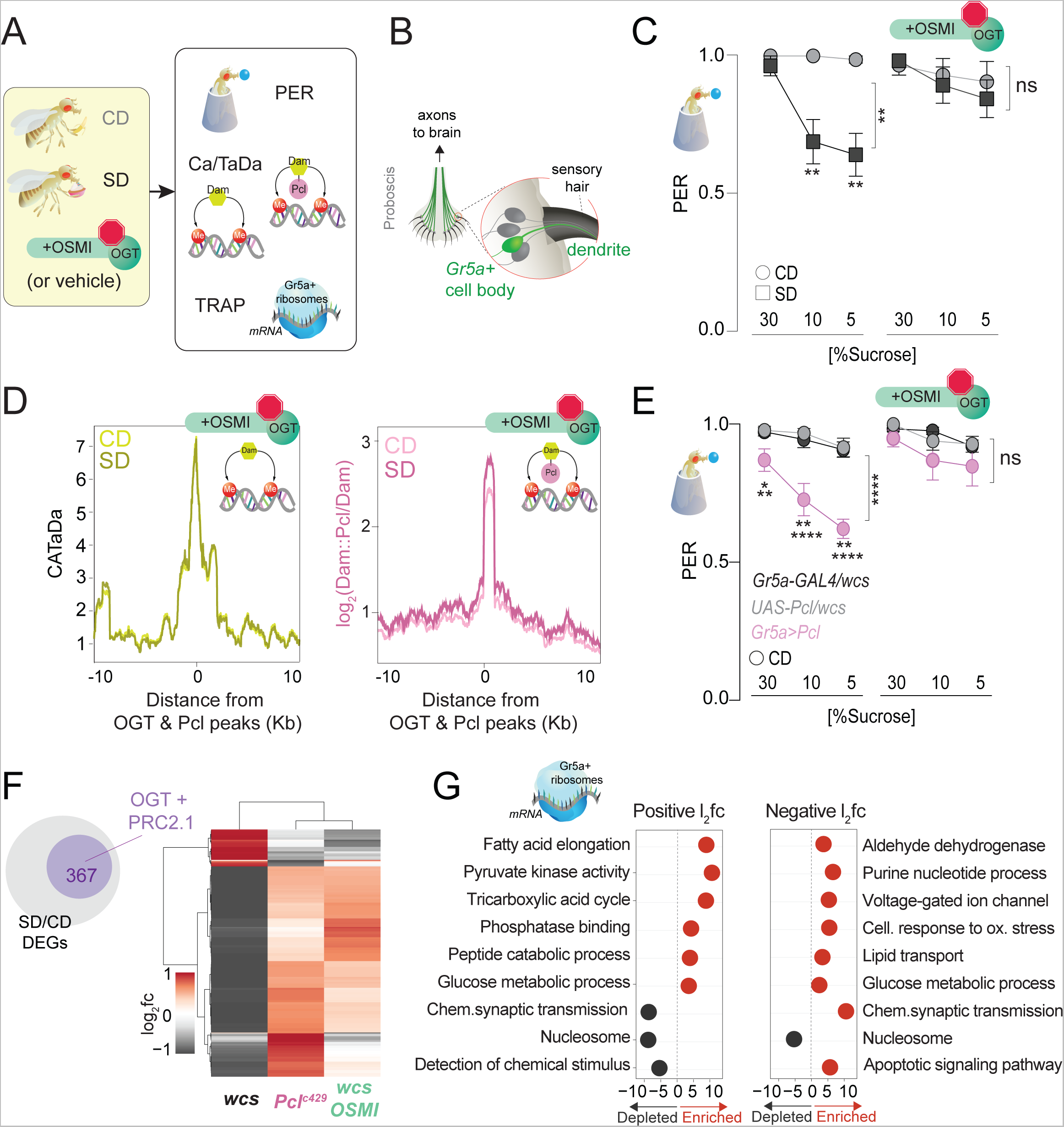
OGT activity is necessary for chromatin, transcriptional, and behavioral dynamics in response to the sugar diet environment. **(A)** Experimental design for the dietary epistasis experiments; control and experimental flies in C-G were exposed to a control or sugar diet with vehicle (DMSO) or the OGT-inhibitor OSMI. **(B)** Diagram of the anatomy of the sensory system showing the cell bodies, dendrites, and axons of the sweet-sensing Gr5a+ neurons. **(C)** Taste responses (*y-axis*) to stimulation of the labellum with 30, 10, and 5% sucrose (*x-axis*) of age-matched male *w1118^cs^*flies on a CD (circle) or SD (square) diet with vehicle (DMSO) or 10μM OSMI. n =14-17. Two way repeated measure ANOVA, main effect of OSMI treatment *p=0.0089*; Tukey multiple comparisons test for 30, 10, and 5% sucrose: 1) CD vs. SD (DMSO) p<0.05, *p=0.0090, p=0.0034* and 2) CD vs. SD (+OSMI), *p>0.05* at all concentrations. **(D) (left)** Average CATaDa signal on a CD (light yellow) and SD (dark yellow) with OSMI-1 centered at *OGT+ Pcl* peaks (compare to Fig. 2D); **(right)** Average log2 Dam::Pcl/Dam signal on a CD (light pink) and SD (dark pink) with OSMI-1 centered at *OGT+ Pcl* peaks (compare to Fig. 2). **(E)** Taste responses (*y-axis*) to stimulation of the labellum with 30, 10, and 5% sucrose (*x* axis) of age-matched male G*r5a>Pcl* (pink) and transgenic controls (shades of gray) on CD supplemented with vehicle (DMSO, n=14-23) or 10 uM OSMI (n=20-42). Two Way Repeated Measure ANOVA: 1) DMSO, main effect of genotype ****(*p<0.0001*) and genotype x concentration (\**p<0.05*); Tukey multiple comparisons test for 30, 10, and 5% sucrose concentrations: *Gr5a>wcs vs. Gr5a>Pcl p=0.0165, p=0.0056, p=0.0025*; *Pcl>wcs vs. Gr5a>wcs,* ns. 2) OSMI: main effect of genotype *p=0.3194* and genotype x concentration *p=0.6893*. **(F)** Log_2_fold (l2fc) of DEGs between SD/CD in *w1118cs* ± OSMI and Pcl^c429^ SD/CD. **(G)** GO term analysis of the DEGs measured in the Gr5a+ neurons of flies fed a CD and SD +OSMI.

We first tested the effects of OGT inhibition on taste plasticity by feeding flies a CD or SD supplemented with 10uM of OSMI or vehicle (DMSO) and conducted the Proboscis Extension Response (PER) assay, which measures taste responses. In this assay, the fly proboscis–which houses the dendrites of the sensory neurons (Fig. 3B)– is stimulated with different concentrations of sucrose. The amount of proboscis extension is measured to generate a taste intensity curve (Fig. 3C). Flies on CD had high PER to both low (5%) and high (30%) sucrose concentrations, that decreased with consumption of a high sugar diet for 7 days (Fig. 3C, left). Inhibition of OGT activity, however, resulted in the same taste responses on both diets (Fig. 3C, right). This effect of OSMI on taste plasticity was identical to that observed with knockdown of OGT in the Gr5a+ neurons (Fig. S3A, green; knockdown efficiency is 50%), suggesting that OGT activity plays an essential role in taste plasticity.

We used this experimental design to then ask how OGT activity affected chromatin accessibility and PRC2.1 binding at OGT x Pcl co-occupied loci in response to diet (Fig. 3A). To do this, we fed *Gr5a>LT3-Dam; tubulin-GAL80* (Fig. 3A) and *Gr5a>LT3-Dam::Pcl; tubulin-Gal80ts* (Fig. S4A) flies a CD or a SD supplemented with OSMI. Strikingly, OSMI treatment completely abolished the changes in chromatin accessibility observed with SD at OGT x Pcl sites (Fig. 3D, compared to Fig. 2D; Supplemental File 1), suggesting that OGT activity is necessary for diet-dependent dynamics at these loci. However, inhibition of OGT activity did not affect Pcl occupancy at these peaks, indicating that recruitment of PRC2.1 to these sites is largely independent of this metabolic enzyme. OSMI also had a mild effect on the occupancy of *Dam::Pcl* genomewide since the number (~1800) and identity (80%) of Dam::Pcl peaks were mainly the same with or without OSMI (Fig. S4C and Fig. S5). Only a smaller fraction of new PRC2.1-only peaks emerged with OSMI treatment (Fig. S4C), and the genes in these intervals were enriched in GO terms such as detection of chemical stimuli, DNA binding, and protein kinase activation (Fig. S5). Thus, OGT activity is required for the diet-dependent decrease in chromatin accessibility but not PRC2.1 recruitment or occupancy. Notably, the catalytic activity of OGT was also necessary for PRC2.1-mediated taste modulation. Overexpression of *Pcl* in the Gr5a+ neurons mimics the effects of SD on taste by decreasing responses to sucrose in flies fed a CD (Fig. 3E, *left*)– a result dependent on the H3K27 methylation activity of this complex (Vaziri et al., 2020). However, OSMI blocked the effects of *Pcl* overexpression on sucrose responses compared to vehicle-fed flies (Fig. 3E, *right*). These results argue for a strong genetic interaction between OGT and PRC2.1 and place this metabolic enzyme upstream of PRC2.1 at both the molecular and behavioral levels.

To understand the consequences of the observed OGT-dependent shifts in chromatin accessibility, we isolated mRNAs associated with the ribosomes of the *Gr5a*+ cells using Translating mRNA Affinity Purification (TRAP) (Chen and Dickman, 2017) in flies fed a CD+OSMI and SD +OSMI. Principal component analysis revealed that most of the variation between samples was due to diet (Fig. S6A); mRNAs specifically expressed in the *Gr5a+* cells, such as the sweet taste receptor genes (*Gr5a*, *Gr64f*, and *Gr64a)* and the fatty acids taste receptor *Ir56D,* were enriched in the *Gr5a+* fraction compared to the input, while bitter receptor genes (*Gr66a* and *Gr32a*) were depleted (Fig. S6B), indicating that the selection of *Gr5a+* mRNAs was successful and comparable to our prior experiments (Vaziri et al., 2020). However, compared to the marked negative skew in gene expression we previously observed with a high sugar diet, where 90% of genes had negative log2 fold changes, OSMI-differentially expressed genes (DEGs) showed a similar distribution in positive and negative changes (Fig. S6C; Supplemental File 1)(Vaziri et al., 2020). Indeed, further analyses revealed that OSMI treatment *reverted* (i.e., showed the opposite direction of change; q<0.1, Wald test) or *restored* (practical equivalence test using a null hypothesis of a change of at least 1.5-fold and q <0.05) the expression of 52% of the DEGs with a SD/CD change (Fig. 3F, gray are downregulated and red are upregulated), and that most of the genes changed with OSMI treatment (367) were also similarly affected by a loss of function mutation in *Pcl* (Fig. 3F). Notable among these were the homeobox *cad* and *Ptx1 and* their regulons, which have been implicated in taste function and plasticity (Fig. S6D) (Vaziri et al., 2020). The genes affected y OSMi treatment were enriched in metabolic and neural processes, such as chemical synapse transmission, synaptic target attraction, cell differentiation, and detection of chemical stimuli (Fig. 3G and S7). These experiments place *OGT* and PRC2.1 in the same genetic pathway that directs diet-induced taste plasticity at the chromatin, transcriptional, and behavioral levels. They also argue that the activity of the metabolic sensory OGT may provide the nutrient-dependent context for PRC2.1-mediated changes in chromatin accessibility.

### Sr is enriched at OGTx PRC2.1 genes and necessary for sweet taste function

Our data show that OGT orchestrates responses to the dietary environment in the sensory neurons. Metabolic flux and the utilization of metabolite pools can be cell-specific, but most, if not all, cells would experience higher OGT activity in response to the high sugar environment (Hart, 2019; Olivier-Van Stichelen et al., 2017). Cell-independent outcomes could arise from the chromatin landscape and PRC2.1 occupancy unique to the Gr5a+ neurons. Still, these do not necessarily provide a specific context to enact changes based on real-time nutritional data. A parsimonious integration between nutrients and cellular context (i.e., neural activity) could occur at the chromatin/transcriptional level. To investigate this possibility, we examined the regulatory regions of OGT x Pcl and OGT x PREs (Polycomb Responsive Elements, DNA motifs to which Polycomb Proteins bind) loci for enriched cis-regulatory motifs (Fig. S8A and B, Supplemental File 1; note that a TF may appear multiple times as dots in this graph because of different binding motifs). In this analysis, the highest log2 fold enrichments were for Polycomb Group proteins like *PhoRC* (1.018, p=0.0099) and Trx-recruiter *Trithorax-like* (*Trl*, 1.972, p=0.0099 and 0.803, p=0.0099; *Trx* is antagonistic to PRC2.1), as well as transcription factors, such as the Zn-finger immediate early gene *Stripe* (*Sr*, homolog of human Early Growth Response 2, EGR2, alias Krox20), the nutrient-sensitive factor *Sterol-Responsive-Element Binding Protein* (SREBP), and the paired domain *pox-neuro* (*poxn*, known to be required for sensory neuron specification) (Supplementary Table 1). To determine if they affected sweet taste, we measured the proboscis extension in response to sucrose when these genes were overexpressed or knocked down in the Gr5a+ neurons. The only factor that affected sweet taste responses across low and high sucrose concentrations was *Sr*, which also showed higher mRNA abundance in the sensory neurons of SD-flies (Fig. S8C).

*Sr* is a conserved transcription factor induced by neural activity via the MAPK/ERK pathway (Beckmann and Wilce, 1997; Chen et al., 2016; Gonzales et al., 2020) and is essential for sensory nerve development and plasticity (Duclot and Kabbaj, 2017; Murphy et al., 1989). This signaling pathway is stimulated by mitogens, such as TGF-β/Activin signaling, which increase with sugar levels, eating, and neural activity in flies and mammals (Lavoie et al., 2020; Liu and Chen, 2022; Wilinski et al., 2019). Of note, OGT x Pcl co-occupied loci were enriched in MAPK/ERK targets (Fig. 2E, blue). Enrichment for Sr cis-regulatory motifs was modest at OGT *or* Pcl loci (l2fc=0.196 and 0.398, respectively; *p<0.001* in both cases via an approximate permutation test) but strong at genes bound by both these factors (OGT x Pcl =0.429 and OGT x PREs=0.976, *p*<0.001, permutation test) (Fig. 4A; Supplemental File 1). When we compared the distribution of Sr sites around the regulatory regions of OGT x PRC2.1 genes (either together or separately), we found a marked bias around the transcriptional start site (TSS) (Fig. 4B, top). In contrast, genes that OGT or PRC2.1 did not occupy, were depleted for Sr binding sites at the TSS (Fig. 4B, bottom). This suggests that Sr predominantly occurs at genes that are bound by OGT *and* PRC2.1. To this end, when we examined the expression of Sr-targets in the Gr5a+ neurons of flies on the two diets, we noticed that these genes had negative log2 fold changes on the sugar diet; this repression, however, was abolished by mutations in *Pcl* and by inhibition of OGT activity (Fig. 4C, compare gray vs. pink and green, respectively). This hints toward functional cooperation between *Sr* and *OGT/PRC2.1*. Importantly, since OGT and PRC2 are not always found at TSS, the bias in *Sr* distribution at the TSS of “OGT/PRC2 regulated genes” indicates that coordination between Sr and OGT/PRC2 may arise not from direct interactions but rather from two distinct paths of information flow.

**Figure 4:**
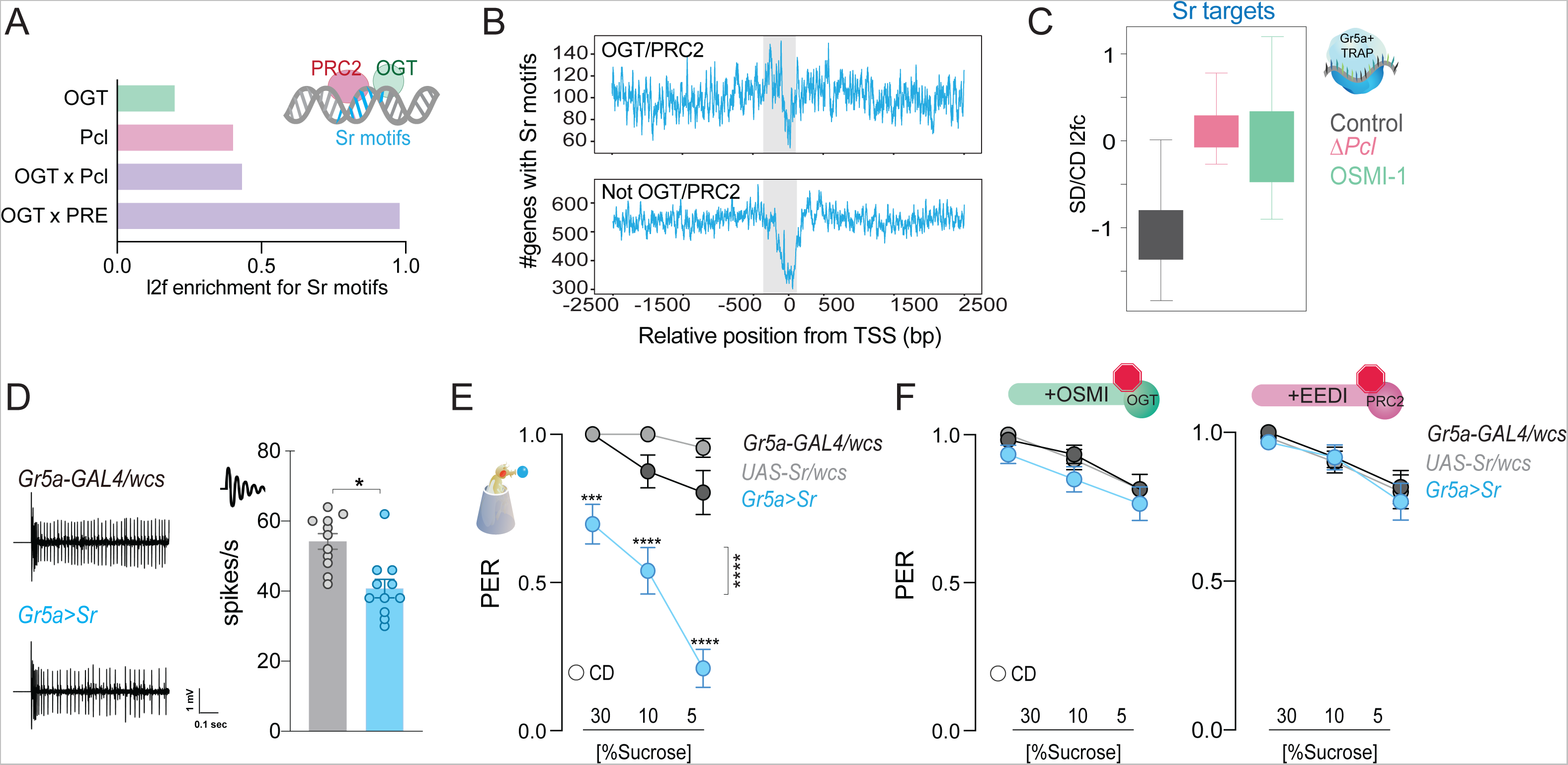
The immediate early gene Sr is found at PRC2 x OGT genes and is involved in sweet taste sensation. (A) Log_2_fold (l2f) enrichment for Sr motifs at sites occupied by OGT (green), Pcl (pink), or OGT+Pcl and OGT+PREs (purple), *p<0.0001*. (B) Distribution of Sr motifs along the regulatory regions 2500 bp up and downstream the TSS for the genes in (A). (C) The distribution of RNA reads normalized as transcripts per million (TPMs) for genes that have Sr sites and are expressed in the Gr5a+ neurons of flies on a CD and SD in control flies or flies with mutations in *Pcl* or inhibition of OGT. *q>0.01*. (D) Representative traces (left) and averaged neuronal responses to 25 mM sucrose stimulation from the L-type sensilla of *Sr* overexpression flies (blue) and controls (gray). n=11. Mann-Whitney test: *p*=0.001. (E) Taste responses (*y-axis*) to stimulation of the labellum with 30, 10, and 5% sucrose (*x-axis*) in *Sr* overexpression flies (blue) and controls (gray). n=22-38. Two way repeated measure ANOVA, main effect of genotype *p<0.0001* and genotype x concentration *p<0.0001*; Tukey post-test for multiple comparisons: 30%: \*\*\**p=0.0002* for *Gr5a>Sr* compared to each control, 10%: *Gr5a>Sr vs. Gr5a>wcs p=0.0025* and *Gr5a>Sr vs. Sr>wcs p<0.0001*; 5%: \*\*\**p*=0.0001 for *Gr5a>Sr* compared to each control. *Gr5a>wcs vs. Sr>wcs p>0.05* at all concentrations. (F) Taste responses (*y-axis*) to stimulation of the labellum with 30, 10, and 5% sucrose (*x-axis*) in *Sr* overexpression flies (blue) and controls (gray) treated with the OGT inhibitor OSMI (green) or the PRC2 inhibitor EEDi (pink). n=30. Two-way repeated measure ANOVA, main effect of genotype *p=0.2993* and *p=0.9146* and genotype x concentration *p=0.9293* and *p=0.9146,* respectively.

To characterize the effects of *Sr* on neural activity and behavior, we used the UAS/GAL4 system to overexpress this gene in the Gr5a+ neurons of adult flies. Overexpression of *Sr* resulted in lower electrophysiological responses of the gustatory neurons to sucrose (Fig. 4D, Mann-Whitney test, *p=0.001*) as well as lower PER at both high and low sucrose concentrations (Fig. 4E). This effect, however, was dependent on the catalytic activities of PRC2.1 and OGT. Indeed, overexpression of *Sr* in flies treated with OSMI (green) or with an inhibitor of PRC2.1 (pink, EEDi, used as in our previous work (Vaziri et al., 2020)) resulted in sucrose responses comparable to those of control flies (Fig. 4F). Overall, these results argue for a role of *Sr* in taste plasticity in coordination with *OGT* and *PRC2.1*.

### The ERK pathway modulates taste adaptations in response to diet

*Sr* is the downstream transcriptional effector for the Extracellular-signal Regulated Kinase (ERK), a pathway stimulated by neural activity that plays a role in neuroplasticity (Lavoie et al., 2020; Miningou and Blackwell, 2020; Thomas and Huganir, 2004) (Fig. 5A). We reasoned that ERK/EGR2 might provide the sensory neurons with a specific context to drive cell-specific outcomes under the same dietary environment. To test this hypothesis, we examined the role of the kinase *rolled* (*rl*)– the ERK homolog in *D. melanogaster*– in sweet taste and diet-induced taste plasticity.

**Figure 5:**
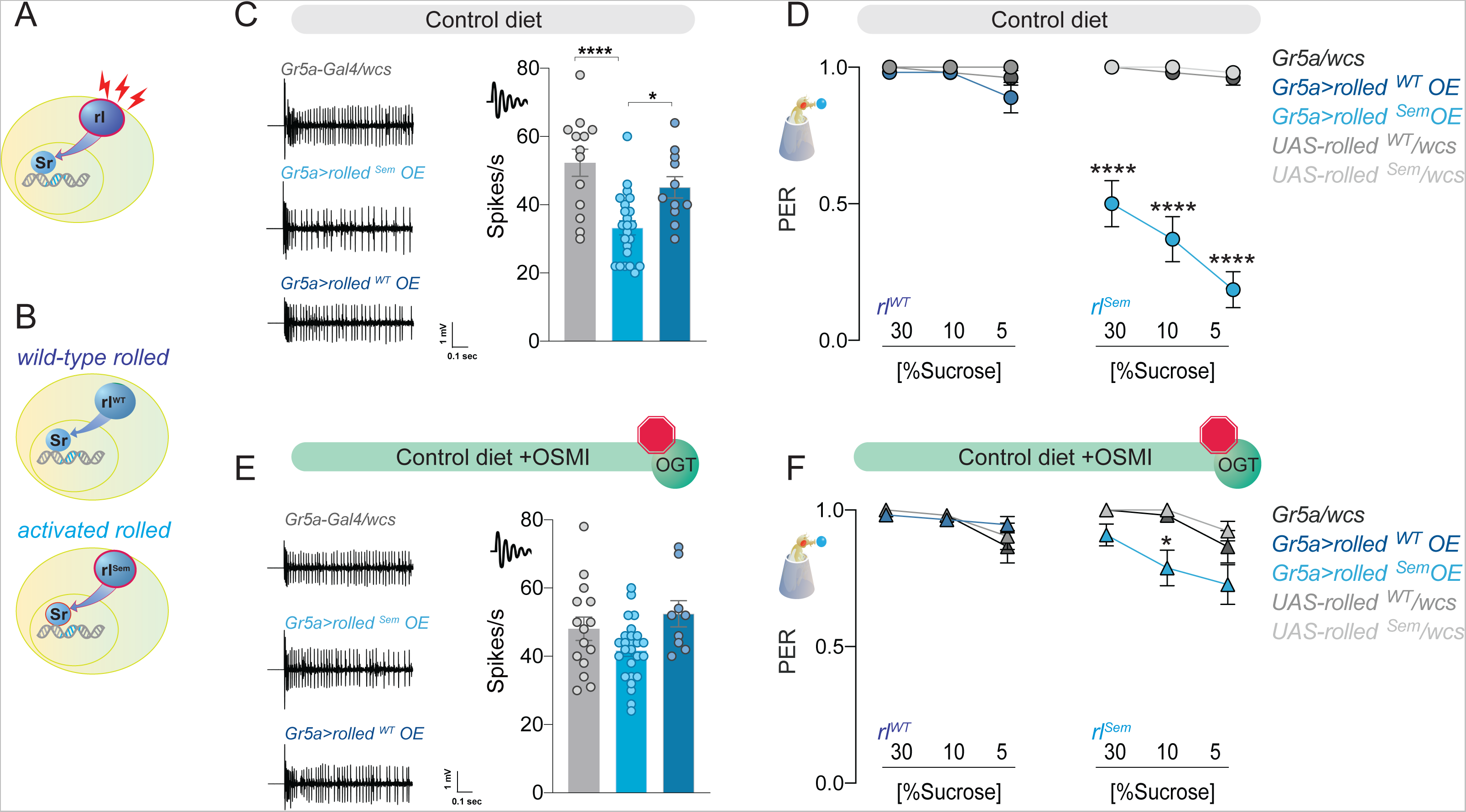
The effect of the kinase rl*/ERK* on sweet taste depends on OGT activity. A. Diagram of the rl/ERK > Sr pathway, red sparks represent neural activity, and red outline represents activation.
B. Diagram of the two types of rl/ERK transgenes used.
C. Representative traces (left) and averaged neuronal responses to 25 mM sucrose of L-type sensilla in overexpression of wild-type (rl^WT^) or constitutively active (rl^Sem^) rl/ERK in the Gr5a+ neurons (blue) and control (gray) on a CD. n=11-23. One way ANOVA; Tukey’s multiple comparison test: *p*<0.0001 for *Gr5a/+* vs. *Gr5a>rl^Sem^*, *p*=0.279 for *Gr5a/+* vs. *Gr5a>rl^WT^*, and *p*=0.018 for *Gr5a>rl^Sem^* vs. *Gr5a>rl^WT^*.
D. Taste responses (*y-axis*) to stimulation of the labellum with 30, 10, and 5% sucrose (*x-axis*) in flies with overexpression of wild-type (*rl^WT^*) or constitutively active (*rl^Sem^*) rl/ERK in the Gr5a+ neurons (blue) and control (gray) flies on a CD+vehicle (DMSO). n=24-27. Two-way repeated measure ANOVA, main effect of genotype *p<0.0001* and concentration x genotype *p<0.0001*. Tukey multiple comparisons tests: *p<0.0001* for *Gr5a>rl^Sem^* vs. all other genotypes at 30, 10, and 5% and *p>0.05* for all other comparisons at all concentrations.
E. Representative traces (left) and averaged neuronal responses to 25 mM sucrose of L-type sensilla in overexpression of wild-type (*rl^WT^*) or constitutively active (*rl^Sem^*) rl/ERK in the Gr5a+ neurons (blue) and control (gray) on a CD+OSMI. n=11-23. One way ANOVA; Tukey’s multiple comparison test: *p*=0.172 for Gr5a/+ vs. *Gr5a>rl^Sem^*, *p*=0.603 for Gr5a/+ vs. *Gr5a>rl^WT^*, and *p*=0.034 for *Gr5a>rl^Sem^* vs. *Gr5a>rl^WT^*
F. Taste responses (*y-axis*) to stimulation of the labellum with 30, 10, and 5% sucrose (*x-axis*) in flies with overexpression of wild-type (*rl^WT^*) or constitutively active (*rl^Sem^*) Rl/ERK in the Gr5a+ neurons (blue) and control (gray) in flies fed a CD+OSMI. n 26=33. Two-way repeated measure ANOVA, main effect of genotype *p=0.005*; Tukey multiple comparisons tests: *p>0.05* for all other comparisons at all concentrations except for *p<0.0001* for *Gr5a>rl^Sem^*vs. *rl^Sem^/wcs* at 10% *p=0.0216.* Effect of OSMI vs. vehicle: *Gr5a>rl^Sem^* 30% *p=0.0012*, 10% *p=0.0030*, 5% *p<0.0001*, and *p<0.05* for all other genotypes.

First, we observed that, as with *Sr*, the mRNA abundance of *rolled* was higher in the Gr5a+ neurons of flies on a SD, but this gene was not a direct target of OGT or PRC2.1 (Fig. S9A). We tried several available antibodies against *rolled* and activated (phosphorylated) *rolled* to establish whether increased transcript levels also resulted in higher activation of this kinase; however, none of them resulted in a reliable signal in our hands. We thus turned to genetic tools to investigate whether higher rolled expression or activity played a role in sweet taste plasticity. To differentiate between these two possibilities, we expressed either a wild-type *rolled* (*rl^WT^*, Fig. 5B top, (Biggs et al., 1994)) or constitutively active form of the kinase (*rl^Sem^* Fig. 5B, bottom) (Oellers and Hafen, 1996) in the Gr5a+ neurons and tested neural and taste responses to sucrose. Overexpression of *rl^WT^* with *Gr5a-GAL4* did not affect the electrophysiological responses of the sensory neurons to sucrose (Fig. 5C, *dark blue*); however, expression of the active *rl^Sem^* decreased neuronal responses to sucrose (Fig. 5C, *light blue*). These activity phenotypes were reflected in the behavioral taste responses to sucrose, with *rl^WT^* flies having identical PER to sucrose as controls and *rl^Sem^*showing reduced PER across high and low sucrose concentrations (Fig. 5D, *left vs. right*; note that controls are shared here, plotted separately for clarity). Thus, *rolled* activity, but not higher levels alone was sufficient to affect sweet taste plasticity. Not surprisingly, given the known function of *rl/ERK* in neural activity, we found that this kinase was also necessary for sweet taste responses, as loss of function mutants for *rolled* had lower electrophysiological and behavioral responses to sucrose (Fig. S9B-C). To test whether there was a synergetic interaction between *rl* and OGT, we repeated the same experiments in the presence of the OGT inhibitor OSMI. Strikingly, OSMI treatment almost entirely blocked the effects of *rl^Sem^* on both neural and behavioral responses to sucrose (Fig. 5E-F).

To characterize the function of the *rl/Sr* pathway on taste plasticity, we used Trametinib, a drug that inhibits ERK activation in animals (currently used for treating melanoma). At concentrations previously used in flies (15.6mM) (Castillo-Quan et al., 2019; Slack et al., 2015), Trametinib blocked the effects of *rl^Sem^* expression on sweet taste responses (Fig. S10A). Treatment with this ERK inhibitor also negated the effects of *Sr* overexpression on sucrose responses, resulting in flies with PER comparable to controls (Fig. S10B). Thus, Trametinib treatment efficiently blocks ERK signaling. To determine if the activity of the *rl/Sr* (ERK/Krox20) pathway was necessary for taste plasticity in response to the sugar diet environment, we fed flies a control or sugar diet with or without Trametinib for 7 days, then measured their neural and behavioral responses to sucrose. Exposure to a high sugar diet decreases the electrophysiological (Fig. 6A) and behavioral (Fig. 6B) responses to sucrose. However, when *rl* activity was blocked with Trametinib, there was no decrease in neural responses or PER (Fig. 6C and 6D). Of note, Trametinib had a minor but significant effect on sweet taste activity (compare CD of Fig. 6C with CD of Fig. 6D), consistent with the observation that *rl* is necessary for normal sweet taste function (Fig. S9B). Together, these data indicate that the ERK pathway plays a critical role in the development of taste adaptations in response to diet and place its function upstream of OGT.

**Figure 6:**
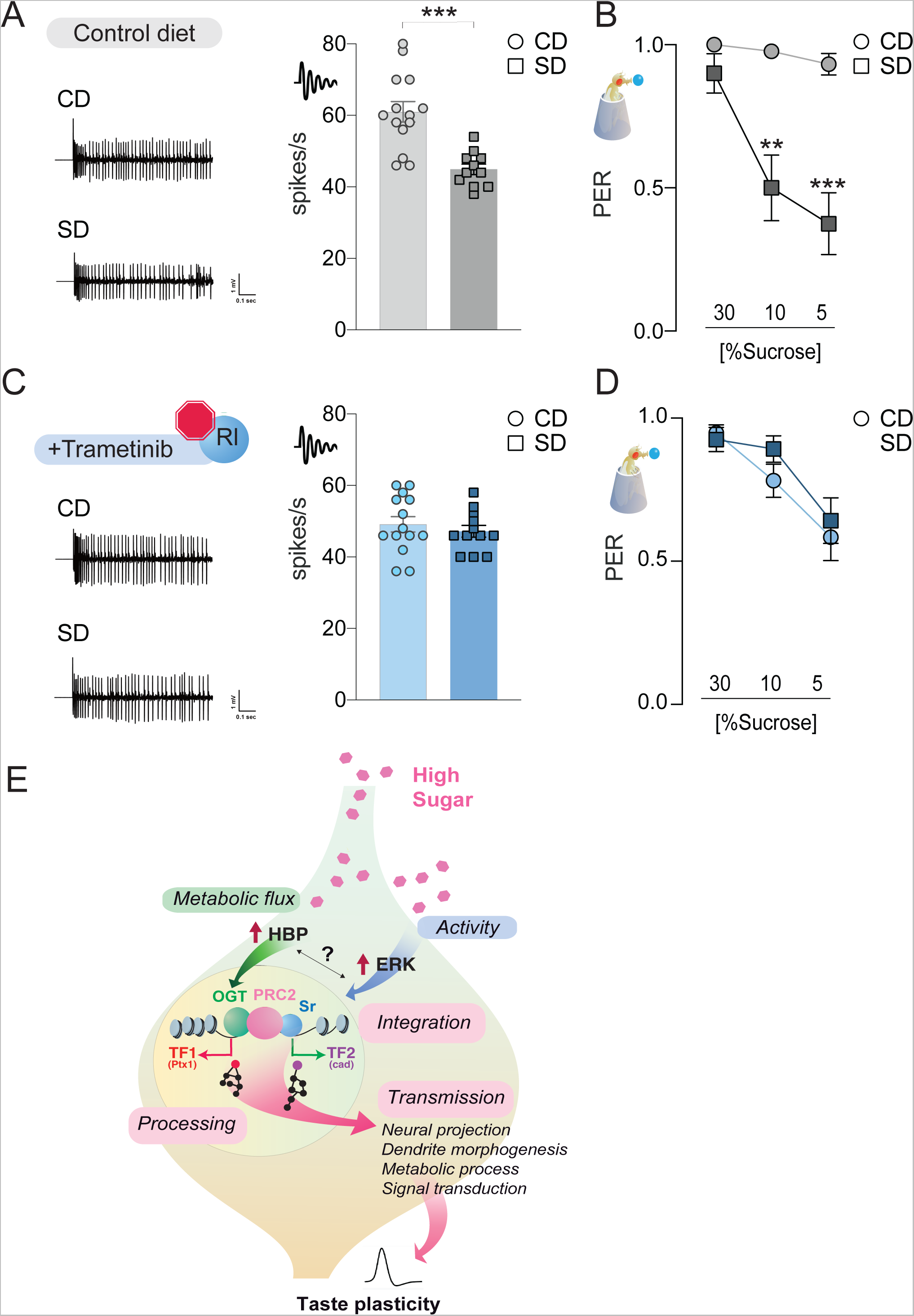
The rl>Sr pathway is important for taste adaptations in response to diet. **A and C)** Representative traces (left) and averaged responses to 25 mM sucrose from L-type sensilla of flies fed a CD and SD (**A**, gray) or Trametinib (**C**, blue). n=11-14. Unpaired t-test: *p*=0.0001 for CD vs. SD, and *p*=0.486 for CD Trametinib vs. SD Trametinib. **B and D)** Taste responses to stimulation of the proboscis with sucrose in flies fed a CD and SD + vehicle (**B**, DMSO, gray) or Trametinib (**D**, blue). PER, n=20-33. PER: Two way repeated measure ANOVA, main effect of diet, vehicle *p<0.0001* and Trametinib *p=0.4701*; Tukey multiple comparison test: vehicle CD vs SD 30% *p=0.421*, 10% *p=0.0017*, and 5% *p=0.0002* and Trametinib CD vs SD 30% *p=0.9702*, 10% p=*0.4470*, and 5% *p=0.9575*. Effect of Trametinib: CD vehicle vs CD Trametinib, 30% *p=0.1745*, 10% *p=0.0108*, 5% *p=0.0015*; SD vehicle vs SD Trametinib, 30% *p=0.4837*, 10% *p=0.0228*, 5% *p= 0.2339*. **E)** A model for how OGT, PRC2, and ERK orchestrate taste plasticity in response to a changing food environment. Boxes in pink describe the different steps of information processing (see discussion).

## DISCUSSION

Nutrigenomic signaling plays a function in health and disease by bridging the dietary environment with physiological adaptations. However, the molecular mechanisms and consequences of this type of nutrient sensing are still poorly understood. In particular, how nutrigenomic signals are integrated with cellular contexts to drive specific outcomes has remained hard to define due to the lack of mechanistic nutrigenomic models (Müller and Kersten, 2003; Vaziri and Dus, 2021). In this work, we exploited the conserved phenomenon of diet-induced taste plasticity and the genetic tools of the *D. melanogaster* fly to answer these questions.

Here we report that the metabolic enzyme OGT is associated with neural chromatin at introns and transcriptional start sites. While OGT-associated genes showed subtle but significant changes in chromatin accessibility in response to diet, these dynamics were much more robust at loci co-occupied by both OGT and the epigenetic silencer PRC2.1. At genes decorated by both factors, we observed sizable diet-dependent chromatin variations that were critically dependent on the catalytic activity of OGT. OGT activity was also necessary for the differential transcriptional and taste responses to the high-sugar diet. The OGT and PRC2 bound nutrient-dependent loci were enriched for binding motifs for the activity-dependent transcription factor Sr, the effector of the ERK pathway. We show that this signaling pathway functions upstream of OGT/PRC2.1 to shape neural and behavioral taste responses to the dietary environment. We thus propose a model where a nutrigenomic pathway composed of OGT and PRC2 integrates information from the nutrient and cellular environment via ERK signaling to orchestrate specific responses to diet (Fig. 6E). This integration occurs at the level of chromatin and modulates the expression of transcription factors and signaling regulators that further amplify and extend the reach of nutrigenomic signaling. Our findings reveal how nutrigenomics contributes to crucial neural plasticity and behavior aspects.

### OGT and Chromatin Dynamics

OGT is a conserved enzyme that catalyzes the transfer of UDP-GlcNAc to the serine and threonine residues of proteins (Hart, 2019; Olivier-Van Stichelen et al., 2017). Because UDP-GlcNAc synthesis by the Hexosamine Biosynthesis Pathway (HBP) combines sugar, amino acid, nucleotide, and fatty acid metabolism, the levels of this metabolite, as well as the activity of the enzymes in this pathway, are inextricably linked to cellular metabolism and diet (Hart, 2019; Olivier-Van Stichelen et al., 2017). Higher HBP flux directly impacts OGT activity because the function of this enzyme is linear across a vast range of physiological UDP-GlcNAc concentrations. Because of this, OGT is recognized as a critical nutrient sensor in animal physiology, particularly in development, cancer, and metabolic disease (Hart, 2019; Olivier-Van Stichelen et al., 2017). More recently, its importance for neural function and plasticity has also been recognized, with studies implicating it in synapse maturation, neural excitability, activity, and plasticity (Ardiel et al., 2018; Butler et al., 2019; Giles et al., 2019; Hwang and Rhim, 2019; Lagerlöf et al., 2017, 2016; Li et al., 2019; May et al., 2019; Ruan et al., 2014; Su and Schwarz, 2017). Our group showed that a high sugar diet acutely and chronically increased HPB activity in flies and played a role in diet-induced sensory plasticity (May et al., 2019; Wilinski et al., 2019).

OGT is a nucleocytoplasmic enzyme, and the GlcNAc modification is enriched in nuclear and synaptic proteins, which extends its reach on cellular physiology (Hart, 2019; Olivier-Van Stichelen et al., 2017). Although OGT is thought to play a role in gene regulation, only one study has shown its direct association with chromatin in murine embryonic stem cells (Vella et al., 2013). Here we show that OGT also decorates neural chromatin in *Drosophila melanogaster*. Similar to embryonic stem cells, OGT was enriched at introns and transcriptional start sites and primarily associated with transcriptionally active chromatin. However, we found that a portion of OGT was also enriched at Polycomb repressive chromatin, consistent with previous reports that the GlcNAc modification is found at PREs, as well as on many PcG proteins; OGT is also associated with PcG complexes to mediate Hox-gene repression (Hart, 2019; Olivier-Van Stichelen et al., 2017; Schuettengruber et al., 2017). On a high-sugar diet, there was a higher association of OGT with DNA but lower chromatin accessibility. However, the magnitude of these changes depended on what other regulatory and DNA-binding factors were found at OGT loci. At loci with PRC2.1 binding and Sr/EGR2 motifs, chromatin openness was markedly reduced in response to the high-sugar diet environment. To the best of our knowledge, this is the first study to show that OGT-associated chromatin is nutrient-sensitive. Importantly, this nutrient sensitivity was entirely dependent on the catalytic activity of OGT because it was abolished in the presence of the inhibitor OSMI-1. Interestingly, OGT activity had no effect on PRC2.1 association with co-occupied loci (and only a small effect at non-OGT loci, data not shown). The H3K27 methylation activity of PRC2.1 is required for changes in chromatin accessibility, including those that depend on diet in the sensory neurons (Schuettengruber et al., 2017; Vaziri et al., 2020). Thus, our data suggest that OGT activity affects the repressive action of PRC2.1.

These findings raise several important questions about the biochemical mechanisms of OGT action that our genetic system is poorly suited to address, but that will be important to define in future studies. First, what are the targets of OGT at nutrient-sensitive loci? Is OGT directly GlcNAcylating PRC2.1 to modify its repressive drive? Several studies have linked OGT activity with the stability, chromatin occupancy, or catalytic function of Polycomb Group Proteins (Hart, 2019; Olivier-Van Stichelen et al., 2017). OGT influences stability and silencing activity of the PRC2 subunit Enhancer of Zeste Homologue 2 (Ezh2) –the enzyme that catalyzes H3K27 methylation; knockdown of OGT decreased H3K27 methylation and Ezh2 occupancy at promoters in cell culture, while OGT overexpression increased it (Chu et al., 2014; Sakabe and Hart, 2010). OGT also affected the binding of Ezh2 to promoters of transcription factors in breast cancer cells, but not its stability (Forma et al., 2018). Depending on cell type and tissue, GlcNAcylation of Ezh2 did not affect its association with PRC2 but decreased its enzymatic activity (Lo et al., 2018) or stability (Jiang et al., 2019; Sui et al., 2020; You et al., 2021). Further, OGT-dependent recruitment of PRC2.1 to *UNC5* family gene promoters led to their silencing in HCT116 colon cancer cells, which has been linked to the worsening of colorectal cancer by a high carbohydrate diet in mammals (Decourcelle et al., 2020). In the hippocampus, knockdown of *OGT* reduced PRC2.1-mediated gene silencing during fear memory consolidation (Butler et al., 2019); GlcNAcylation of Polyhomeotic– motifs which are enriched at OGT x Pcl sites in our data– was also required for its repressive properties (Gambetta et al., 2009; Gambetta and Müller, 2014). Thus, converging evidence suggests that OGT impacts different aspects of PRC2 and PcG biology and is broadly consistent with our data. Connections between OGT and ERK have also been uncovered in the context of cancer and cell division, with studies showing that inhibition of ERK signaling decreases O-GlcNAcylation and vice-versa (Jiang et al., 2016; Zhang et al., 2015) and that GlcNAcylation promotes ERK-effects while OGT inhibition blocks them (Cork et al., 2018; Lei et al., 2020; Weiss et al., 2021). These findings are consistent with the effect and direction of the genetic interactions we observed between OGT and PRC2.1 and OGT and ERK, the direction of “information flow” within the cell (Fig. 6E), and the effects of our genetic manipulations. OGT could also affect chromatin accessibility by GlcNAcylating histones, although the function and effects of these histone modifications are still poorly understood (Gambetta and Müller, 2015; Hirosawa et al., 2018; Konzman et al., 2020; Olivier-Van Stichelen et al., 2017). Another outstanding question is how OGT is targeted to chromatin and whether this recruitment is dynamic and related to nutrient availability. For example, are there different local pools of OGT and GlcNAc in the nucleus vs. cytoplasm (or mitochondria) where OGT has been described? Finding answers related to cellular compartmentalization of this metabolic enzyme and its targets will be an essential step in understanding nutrient signaling, currently one of the central questions in the field.

### Sensors and effector mechanisms of nutrigenomic signaling

OGT provides an instructional role in cellular adaptations to the dietary environment, as our data and that of others suggest (Hart, 2019; Olivier-Van Stichelen et al., 2017). But OGT is active in every cell, so what determines the specificity of outcomes? The first mode of specificity is likely permissive and related to the general accessibility of genomic loci to change: those silenced during development would also be shielded from the effects of environmental variation.

However, prescriptive mechanisms should also exist. In the case of the sweet taste neurons, sugar directly activates the cells via receptor-dependent mechanisms and enhances OGT’s metabolic activity. Integration of these two signals– a synergy almost entirely unique to these cells– could shape cell-specific outcomes. Our data support this idea by revealing a function for the mitogen-activated protein kinase ERK in neural and behavioral taste adaptations. *rl/ERK* and its effector, *Sr/EGR2* mRNAs, were higher in the Gr5a+ neurons of flies on a sugar diet. Binding sites for the ERK effector Sr/EGR2 are among the most enriched at OGT x Pcl/PRE loci, and several of these co-occupied genes are known to be regulated by ERK, like the transcription factor c-fos (*kayak*).

Further evidence of ERK/EGR2 interactions with OGT/PRC2 comes from the observations that diet-driven changes in Sr/EGR2 regulons, as well as the effects of Sr overexpression on sweet taste, depended on the activity or presence of OGT and PRC2. Most importantly, we demonstrate that this pathway affects taste plasticity at both the neurophysiological and behavioral levels, with strong epistatic interactions with OGT. Together, these molecular and functional data support the idea that the activity-dependent ERK pathway provides the context (likely neural activity) for cell-specific outcomes in response to the same nutrient environment. It is worth noting that this pathway has the potential to tune plasticity to other nutrient-activity fluctuations, such as those experienced with fasting and satiety, since flux through the HBP and known-ERK activators, such as TGF-β, change with acute and chronic energy changes (Hart, 2019; Lavoie et al., 2020; Liu and Chen, 2022; Olivier-Van Stichelen et al., 2017; Papa et al., 2019; Wilinski et al., 2019).

Finally, sensing mechanisms must be transformed into actions to be effective. This is likely the role of PRC2.1. Only a small portion of the genes occupied by PRC2.1 is sensitive to diet and OGT activity. PRC2.1 decorates chromatin at these sites already in the control diet condition but redistributes (up or down) in response to the high sugar diet (Vaziri et al., 2020). Thus, PRC2.1 is not binding to new loci but instead tuning the output of those it is already bound to.

Further, OGT activity did not affect PRC2.1 occupancy at OGT x Pcl sites, suggesting that other mechanisms mediate this effect. To this end, the Tudor domain of the mammalian orthologues of Pcl, MTF2, binds H3K36m to promote the intrusion of PRC2.1 into active genes and drive repression (Ballaré et al., 2012; Brien et al., 2012; Cai et al., 2013). This could be a potential mechanism through which PRC2.1 is redistributed to transcribed loci in the sugar diet environment. Perhaps activity- or nutrient-transcription could play a role in this process. What mechanisms promote PRC2.1 activity or nutrient-dependent redistribution will be important to define in the future but will be better addressed in vitro models.

### From nutrigenomics to neural plasticity

In our model, metabolic and activity sensors integrate cellular information to promote changes in gene expression. How are these actualized into physiological (in this case, neural) adaptations that underlie behavior or disease? This is one of the central and unresolved questions in nutrigenomics. Our model’s genetic and neural tractability provides a unique opportunity to get some answers. The 162 co-occupied loci identified were enriched for transcription and regulatory factors involved in cell proliferation, differentiation, signaling, and neural activity, as well as pathways implicated in neural plasticity. Many of these DNA binding factors play essential roles during the development of sensory neurons to set their biophysical properties, such as *Ptx1* and *cad*, but also affect adult taste plasticity. The regulons of these TFs include genes known to affect pre and postsynaptic branching and structure, as well as synaptic physiology. Thus our collective data indicate that this nutrigenomic pathway promotes taste adaptations most likely by re-engaging developmental gene batteries, a mechanism that has been hypothesized to play a role in neural plasticity (Hobert, 2011; Marder and Prinz, 2002; Parrish et al., 2014; Vaziri et al., 2020). Whether this is a general rule of nutrigenomic signaling or something specific to these sensory neurons is yet to be determined; however, it is interesting to note that this is similar to how cancer cells exploit developmental networks for uncontrolled growth (DeBerardinis and Chandel, 2016; Faubert et al., 2020).

## LIMITATIONS

Although using sensory plasticity and fly gustatory neurons as a model to study nutrigenomic signaling brings unique advantages, it also has significant limitations. These primarily arise from the small number of cells (60) and the *in vivo* nature of our model. First, we cannot probe whether OGT, PRC2, and EGR2 physically interact or modify each other in these cells. Thus, evidence for our model arises from the combination of cell-specific molecular, genetic, and physiological data. Second, we only inferred that the loci with Sr/EGR2 motifs integrate activity due to the well-established function of the ERK pathway in activity-dependent plasticity; future studies should address this directly and compare the effects of acute vs. chronic nutrient influx.

Further, while inhibitors have allowed us to establish critical epistatic interactions and conduct dietary manipulations while bypassing developmental effects and genetic challenges, we cannot exclude that some of these effects may be non-cell autonomous. Integrating this model with biochemical approaches that preserve the appropriate activity and nutrient context would help address these critical questions. Finally, pathways beyond OGT, ERK, and PRC2.1 may also play a role in sensory plasticity.

## CONCLUSIONS

In summary, we show that activity and nutrient sensing mechanisms are integrated at the genomic level to promote neural adaptations to the food environment. In particular, our data reveals a central and instructional role for OGT and meaningful epistatic interactions with sensors (ERK) and effectors (PRC2.1). More generally, we put forth a model where cell and context specificity transforms “nutritional data” – i.e., variations in nutrient and metabolite levels– into nutritional information (Floridi, 2005), as shown in Fig. 6E (pink boxes). This information is processed and interpreted by gene regulatory processes to make “decisions’’ about responding to environmental challenges and carrying out physiological, neural, and behavioral changes. Thus, nutrigenomic mechanisms could provide a critical path for information flow in biological systems (Fabris, 2009; Reinagel, 2000; Shannon, 1948; Smith, 2000). A clear advantage could reside in their ability to amplify transient, and often minor, variations in nutrient and activity levels into strong reactions, which can be used to orchestrate responses to current *and* future environmental challenges. Future studies in this field will no doubt uncover fascinating insights about the rules of nutrigenomic communication: these discoveries will illuminate how nutrition and gene expression converge to shape cell physiology and provide us with new tools to promote wellness and diminish the burden of disease.

## METHODS

### Fly husbandry, strains, and diets

All flies were grown and fed cornmeal food (Bloomington Food B recipe) at 25°C and 45 to 55% humidity under a 12-hour light/12-hour dark cycle (Zeitgeber time 0 at 9:00 AM.) unless otherwise stated. Male flies were collected under CO_2_ anesthesia 1-3 days after eclosion and maintained in a vial that housed 35-40 flies. Flies were acclimated to their new vial environment for an additional 2 days and were moved to fresh food vials every other day. The GAL4/UAS system was used to express transgenes of interest using the *Gustatory receptor 5a Gr5a-GAL4* transgene. For each GAL4/UAS cross, transgenic controls were made by crossing the *w1118^CS^* (gift from A. Simon, *CS* and *w1118* lines from the Benzer laboratory) to GAL4 or UAS flies, sex-matched to those used in the GAL4/UAS cross. The fly lines used for this paper are listed in supplemental file 1.

For all dietary manipulations, the following compounds were mixed into standard cornmeal food (Bloomington Food B recipe) (0.58 calories/g) by melting, mixing, and pouring new vials as in (Musselman and Kühnlein, 2018) and (Na et al., 2013). For the 30% sugar diet (1.41 calories/g), Domino granulated sugar (w/v) was added. Inhibitors were solubilized in 10% DMSO and added to the control o sugar diet at a total concentration of 10 μM for OSMI (May et al., 2020; Ortiz-Meoz et al., 2015), 8 μM for EEDi (Vaziri et al., 2020), and 15.6mM for Trametinib (Castillo-Quan et al., 2019; Slack et al., 2015). Animals were assigned randomly to dietary groups. The sample sizes were determined based on standards in the field. No animal was excluded from any of the analyses.

### Proboscis extension response

Male flies were food-deprived for 18 to 20 hours in a vial with a Kimwipe dampened with Milli-Q filtered deionized water. Proboscis Extension Response was carried out as described in (Shiraiwa and Carlson, 2007). Scoring of extension responses was conducted manually, and whenever possible, experimenters were blinded. Experiments were replicated 2-3 times by two different experimenters.

### Affinity purification of ribosome-associated mRNA (TRAP)

Male fly heads (300 per replicate, ~10,000 *Gr5a+* cells) were collected using sieves chilled in liquid nitrogen and dry ice. Frozen tissue was then lysed as previously described (Chen and Dickman, 2017; Vaziri et al., 2020). From 10% of the total lysate, total RNA was extracted by TRIzol LS Reagent (Thermo Fisher Scientific, 10296010) for input. The remainder of the lysate was precleared by incubation with Dynabeads Protein G (Thermo Fisher Scientific, 10004D) for 2 hours and subsequently incubated with Dynabeads Protein G and an anti-Flag antibody (Sigma-Aldrich, F1804) at 4°C with rotation for 2 hours, then RNA was extracted from ribosomes bound to beads by TRIzol Reagent (Chen and Dickman, 2017).

### Targeted DNA adenine methyltransferase identification (TaDa) and Chromatin Accessibility TaDa (CATada)

To generate the *UAS-LT3-Dam::OGT* construct, the coding region of the *OGT* gene was amplified from *w1118^CS^* animals with the primers listed below and assembled into the *UAS-LT3-DAM* plasmid (gift from A. Brand, University of Cambridge) using the NEBuilder HiFi DNA Assembly kit based on the manufacturer’s instructions (New England Biolabs). Transgenic animals were validated by reverse transcription PCR targeting the correct insert. The *UAS-LT3-Dam::OGT* and *UAS-LT3-Dam* line were crossed to the *Gr5a-GAL4*; *tubulin-GAL80^ts^*. All animals were raised and maintained at 20°C. Expression of *Dam::OGT* and *Dam* was induced at 28°C for 18 hours. For all experiments, 300 heads of male and female flies were collected per replicate on dry ice by sieving. DNA was extracted following kit instructions (Invitrogen). To identify methylated regions, purified DNA was digested by Dpn I, followed by PCR purification of digested sequences. TaDa adaptors were ligated by T4 DNA ligase (NEB). Adapter ligated DNA was PCR-amplified and purified according to the protocol (Marshall et al., 2016). Purified DNA was digested with Dpn II, followed by sonication to yield fragments averaging 300 base pairs. TaDa adaptors were removed from sonicated DNA by digestion followed by PCR purification, and purified sonicated DNA was used for library preparation (Vaziri et al., 2020) and (Marshall et al., 2016).

pUAST-Sxc.Forward gatctgGCCGGCGCaATGCATGTTGAACAAACACGAATAAATATG

pUAST-Sxc.Reverse gttccttcacaaagatcctTTATACTGCTGAAATGTGGTCCGGAAG

### Library preparation

Generation of RNA sequencing (RNA-seq) libraries was with the Ovation SoLo RNA-seq System for *Drosophila* (NUGEN, 0502-96). All reactions included integrated Heat-Labile Double-Strand Specific DNase treatment (ArcticZymes, catalog no. 70800-201). The DNA-sequencing libraries for TaDa were generated using the Takara ThruPLEX Kit (catalog no. 022818). For rat RNA-seq, libraries were prepared using the Nugen Ovation Model organism (Rat #0349-32) with 1/10th ERCC spike-in mix. These libraries were run on a NextSeq instrument using a HO 150 cycle cit (75×75 bp paired-end reads). All *Drosophila* libraries were sequenced on the Illumina NextSeq platform (High-output kit v2 75 cycles) at the University of Michigan Genomics Core facility.

### High-throughput RNA-seq analysis

Fastq files were assessed for quality using FastQC (Andrews and Others, 2010). Reads with a quality score below 30 were discarded. Sequencing reads were aligned by STAR (Dobin et al., 2013) to dmel-all-chromosomes of the dm6 genome downloaded from Ensembl genomes. Counting was conducted by HTSeq (Anders et al., 2015). Gene counts were used to call differential RNA abundance by DESeq2 (Love et al., 2014). A pipeline was generated from (Wilinski et al., 2019). To determine the efficiency and cell specificity of the TRAP, pairwise comparisons were made between the *Gr5a+*-specific fraction and the input. For comparisons between dietary conditions, DESeq2 was only applied to the *Gr5a+*-specific IP condition. SD7 and *Pcl^c429^* datasets were analyzed from and described in (Vaziri et al., 2020). A cutoff of *q* < 0.1 was used to call DEGs. To identify overlap between datasets GeneOverlap was used (Shen and Sinai, 2013).

### High-throughput TaDa and CATaDa analysis

Fastq files were assessed for quality using FastQC (Andrews and Others, 2010). Reads with a quality score below 30 were discarded. The damidseq_pipeline was used to align, extend, and generate log2 ratio files (*Dam::Sxc/Dam*) in GATC resolution as described previously (Marshall and Brand, 2015) and as in (Vaziri et al., 2020). Reads were mapped by Bowtie2 (Langmead and Salzberg, 2012) to dmel-all-chromosomes of the dm6 genome downloaded from Ensembl genomes, followed by read extension to 300 bp (or to the closest GATC, whichever is first). Bam output is used to generate the ratio file in bedgraph format. Bedgraph files were converted to bigwig and visualized in the UCSC Genome Browser. Principal components analysis plots between biological replicates were computed by multibigwigSummary and plotCorrelation in deepTools (Ramírez et al., 2016). Peaks were identified from ratio files using find_peaks (FDR <0.01) (Marshall and Brand, 2015) and as in (Vaziri et al., 2020). Overlapping intervals or nearby intervals (up to 50bp) were merged into a single interval using mergeBed in BEDtools (Quinlan and Hall, 2010). Intervals common in at least 2 replicate peak files were identified by Multiple Intersect in BEDtools and used to generate the consensus peaks (Quinlan and Hall, 2010). For CATaDa experiments, all analyses were performed similarly to those of TaDa with the exception that *Dam* only profiles were not normalized as ratios but shown as normalized binding profiles generated by converting bam files to bigwig files normalized to 1× dm6 genome as reads per genome coverage (Sequencing depth is defined as the total number of mapped reads times the fragment length divided by the effective genome size). Binding intensity metaplots were made by computing a matrix for specified regions (Ramírez et al., 2016). To determine the proportion of genes that fit within the various chromatin domain subtypes, we first matched *Dam::OGT/Dam* targets to coordinates identified by (Filion et al., 2010) and then determined their gene count in each chromatin subtype (observed) compared to the whole genome (expected). Peak annotations were conducted using the HOMER annotatePeaks tool (Heinz et al., 2010) with the dm6 reference genome. In TaDa analysis, genes were considered targets of the factor being investigated if a peak existed anywhere on their length.

### Pathway enrichment analysis

For all fly experiments, GO term enrichment analysis was performed using the iPAGE package (Goodarzi et al., 2009), using gene-GO term associations extracted from the Flybase dmel 6.08 2015_05 release. For all analyses, iPAGE was run in discrete mode. Independence filtering was deactivated for all discrete calculations. All other iPAGE settings default values. All shown GO terms pass the significance tests for overall information described in (Goodarzi et al., 2009). For each term, bins that are outlined show especially strong contributions [*P* values such that a Benjamini-Hochberg FDR (Benjamini and Hochberg, 1995) calculated across that row yields *q* < 0.05].

### Analysis of cis-regulatory enrichments

For each *D. melanogaster* DNA binding protein motif available from the CIS-BP database (Weirauch et al., 2014), we scanned the *D. melanogaster* genome (dmel 6.08 2015_05 release) using the FIMO binding site discovery tool [cite :doi:10.1093/bioinformatics/btr614]. Hits for each motif were retained as potential binding sites and used to calculate overlaps with other features (e.g., OGT of Pcl sites), as noted. Permutation tests to assess significance were performed through repeated application of the bedtools shuffle [cite: doi.org/10.1093/bioinformatics/btq033] command to obtain 100 resamplings (Supplementary Table 1) or 1,000 resamplings (Figure 4) of the feature location of interest, requiring non-overlap of the randomly placed features. For the analysis in Supplementary Table 1, we separately considered each potential motif for each transcription factor extracted from the CIS-BP database (separate motifs for the same factor are denoted by the gene name followed by a “_#” suffix, with # and integer). In the case of analysis of Sr binding sites in Figure 4, we obtained a merged set of potential Sr binding sites by filtering potential binding sites at a q value threshold of 0.1 (acting separately for each motif) and combining all of the locations that were counted as a potential binding site for any of the Sr motifs available from CIS-BP. Enrichments of overlaps with OGT, Pcl, and PRE sites were calculated by comparing the actual observed count of overlapping features with the mean overlap observed across 1,000 random samplings of the Sr motif locations (preserving the chromosome on which each motif is located during shuffling). For comparison of Sr motif locations with TSSs, we first identified the (strandedness-aware) start location of all “gene,” “mobile_genetic_element,” or “pseudogene” features from the dmel6 Genbank annotations and then categorized all of these locations as “OGT/PRC2” or “Not OGT/PRC2” based on whether or not the gene was associated with an OGT, Pcl, or PRE location (see Supplementary Tables for gene lists for each feature type). The density of Sr motif hits (as defined above) was then calculated as a function of position relative to the TSS.

### Electrophysiology

Extracellular recording on labellar sensilla was performed using the tip recording method (Delventhal et al., 2014). 10-13-day-old files were anesthezied by short ice exposure. To immobilize the proboscis, the reference electrode containing the Beadle-Ephrussi Ringer solution was inserted through the thorax into the labellum. The neuronal firing in L-type sensilla was recorded with a recording electrode (10-20 μm dismeter) containing 25 mM sucrose dissolved in 30 mM tricholine citrate as electrolyte. The recording electrode was connected to TastePROBE (Syntech, The Netherlands), and electrical signals were obtained using the IDAC acquisition controller (Syntech). The signals were amplified (10x), band-pass-filtered (100-3000 Hz), and sampled at 12 kHz. Neuronal firing rates were analyzed by counting the number of spikes for a 500 ms period starting from 200 ms after contact using the Autospike 3.0 software. Experimenters were blinded in the initial characterization of the phenotypes and experiments were independently performed at least 3 times.

### Data analysis and statistics

Statistical tests, sample size, and *p* or *q* values are listed in each figure legend. One or two-way repeated measure ANOVA with posthoc tests were used for all PER experiments. All behavioral data were tested for normality, and the appropriate statistical tests were applied if data were not normally distributed. For the RNA-seq expression datasets, we coupled our standard differential expression with a test for whether each gene could be flagged as “significantly not different” -- i.e., a gene for which we can confidently state that no substantial change in expression occurred (rather than just a lack of evidence for change, as would be inferred from a large p-value on the differential expression test). Defining a region of practical equivalence as a change of no more than 1.5-fold in either direction, we tested the null hypothesis of a change larger than 1.5-fold using the gene-wise estimates of the SE in log2fold change (reported by Deseq2) and the assumption that the actual l2fcs are normally distributed. Rejection of the null hypothesis is evidence that the gene’s expression is not changed substantially between the conditions of interest. Python code for the practical equivalence test can be found on GitHub as calc_sig_unchanged.py. All data in the figures are shown as means ± SEM, **** *P* < 0.0001, *** *P* < 0.001, ** *P* < 0.01, and \**P* < 0.05, unless otherwise indicated.

### Data and material availability statement

All high-throughput data are available at the GEO repository: GSE188757 and GSE146245. LT3-Dam::OGT flies are available upon request; all other fly lines are available in the BDSC database.

### Contributions

Experiments and data analysis: AV, HS, MD, DW, PF

Project design: MD, AV, HS

Data Interpretation: MD, AV, HS, DW, PF

Manuscript writing: MD

Manuscript editing: MD, AV, HS, DW, PF

Project supervision and funding: MD

## Acknowledgments

We thank the University of Indiana at Bloomington, the VDRC, the FLYORF stock collections, and all the investigators who shared fly lines with us. Julia Kuhn designed some of the graphics for the manuscript. We are grateful to Dr. Morteza Khabiri for assistance in the calculation of potential TF binding sites.

## Funding

This work was funded by NIH R00 DK-97141 and NIH 1DP2DK-113750, the Klingenstein-Simons Fellowship in the Neurosciences, the Rita Allen Foundation, and NSF CAREER 1941822 (all to MD), the Rackham Predoctoral Fellowship (to AV), NIH T32 DA007268 (DW), NIH P30 DK089503 (MD and DW), and NIH R35GM128637 (to PLF).

## Conflict of interest

The authors declare no conflict of interest.

## SUPPLEMENTARY FIGURES & LEGENDS

**Figure S1-.**
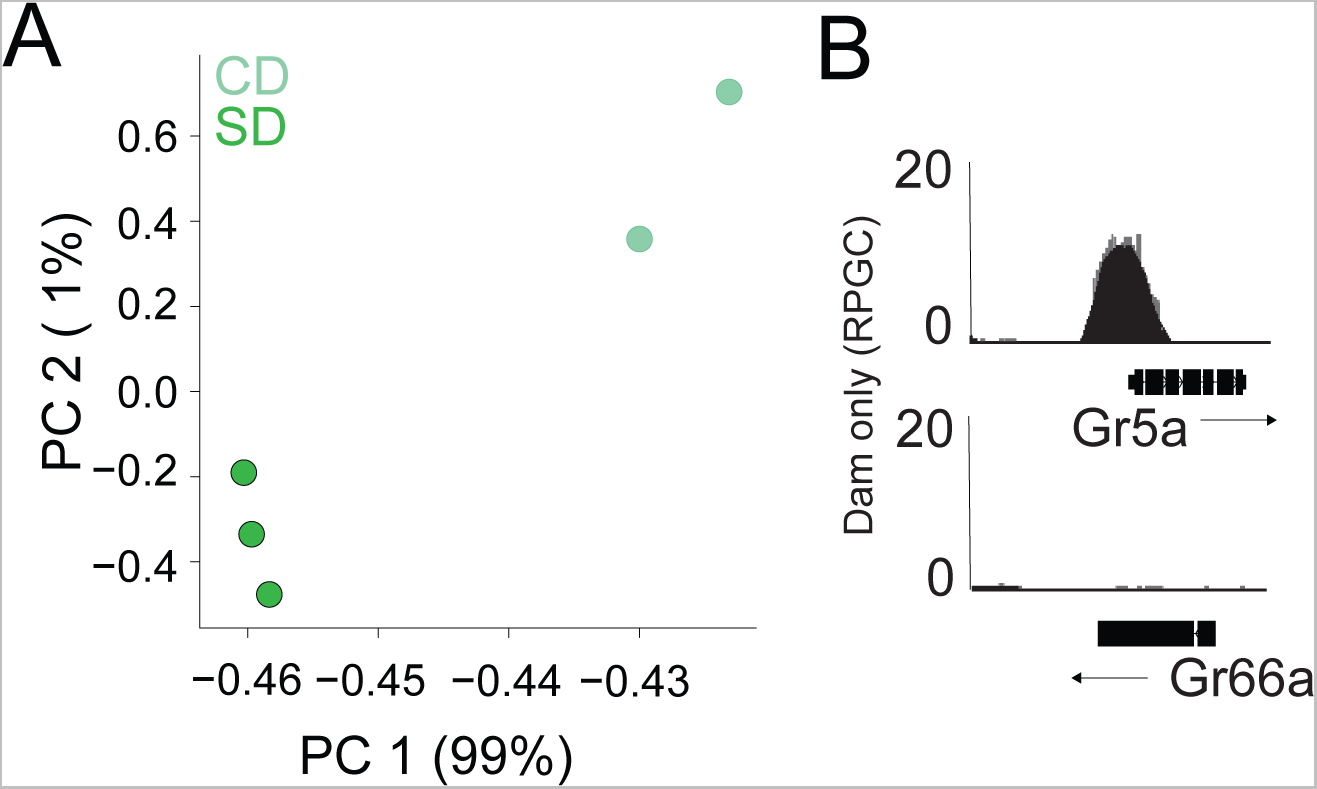
Related to Figure 1, OGT resides on the chromatin of the *Gr5a+* neurons. **(A)** Principal component analysis of normalized log2(*OGT::Dam/Dam*) flies on CD (light green) or SD (dark green). **(B)** CATaDa from control diet flies at the sweet gustatory receptor *Gr5a* and the bitter gustatory receptor *Gr66a*.

**Figure S2-.**
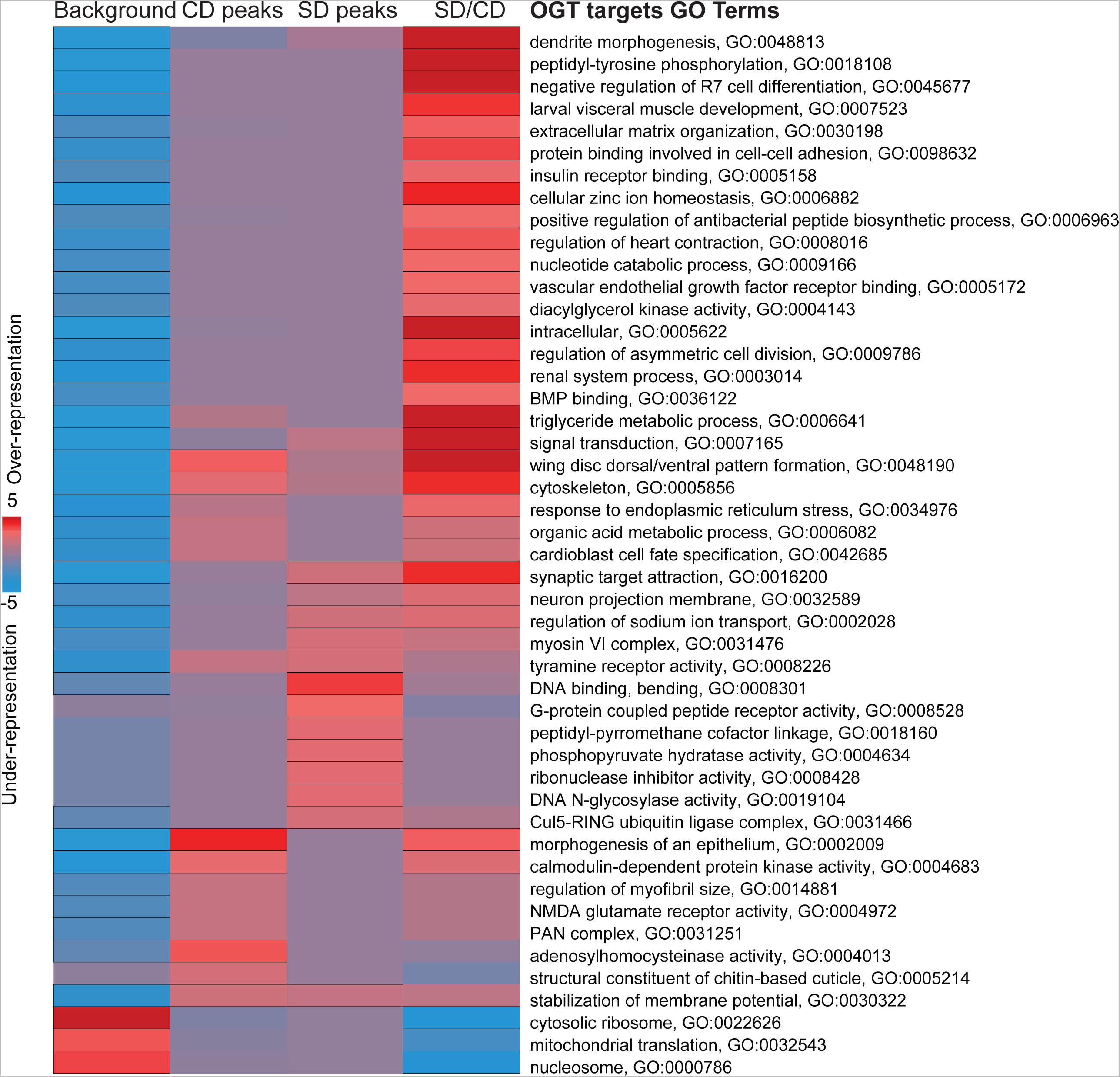
Related to Figure 1, Pathway enrichment analysis of OGT chromatin targets in the *Gr5a+* neurons. iPAGE identification of pathways depleted (blue) or enriched (red) compared to background gene list from the *OGT::Dam* peaks on a control, sugar, and difference of sugar/control diet. The scale represents over-representation (red) or under-representation (blue) of genes within a specific bin for the corresponding GO term. Black outlined boxes represent *q<0.05*.

**Figure S3-.**
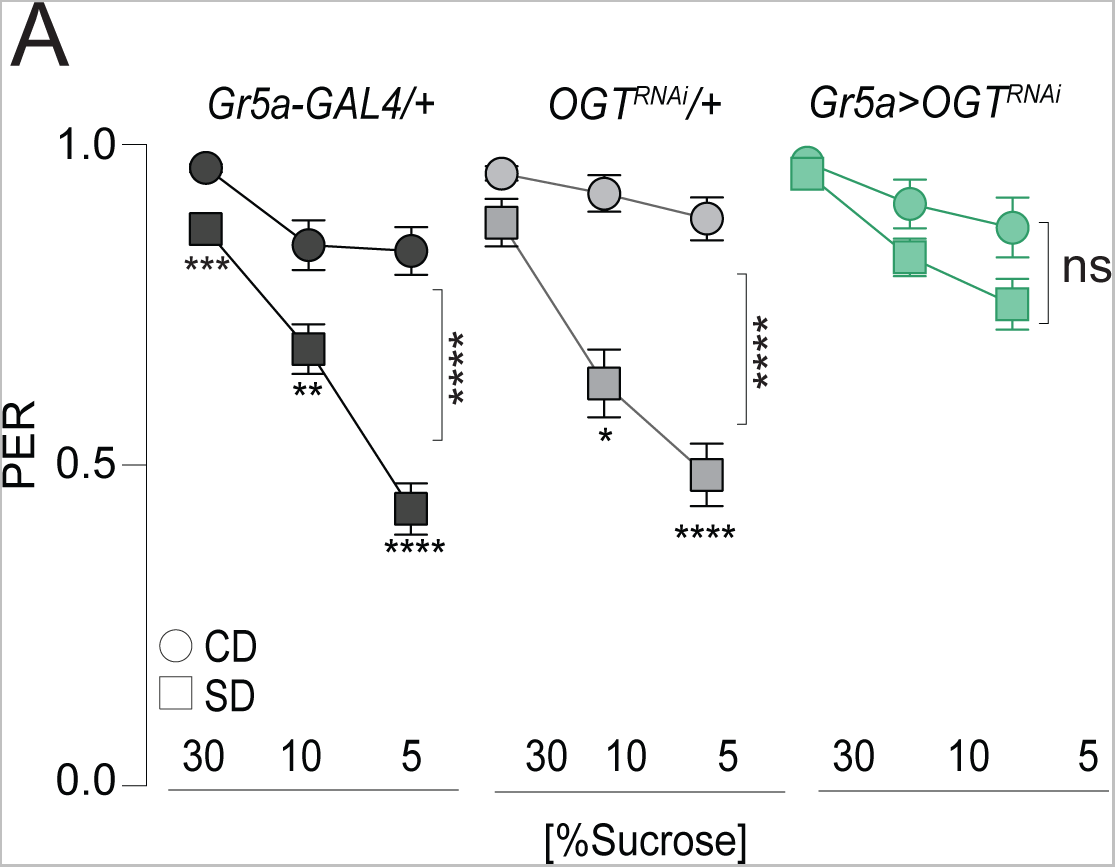
Related to Figure 3, OSMI treatment phenocopies OGT KD in the Gr5a+ neurons. Taste responses (*y-axis*) to stimulation of the labellum with 30, 10, and 5% sucrose (*x-axis*) in flies with knockdown of OGT (green) or controls (shades of gray) in flies fed a CD (circles) or SD (squares). n=18-51. Two way repeated measure ANOVA, main effect of genotype: *Gr5a>wcs p<0.0001* (Tukey multiple comparison 30% *p=0.0008*, 10% *p=0.0047*, 5% *p<0.0001*), *Gr5a>OGT-RNAi p=0.2657* (Sidak multiple comparison 30% *p=0.2792*, 10% *p=0.9756*, 5% *p=0.4883*), *OGT-RNAi>wcs* (Sidak multiple comparison 30% *p=0.5923*, 10% *p=0.0381*, 5% *p<0.0001*).

**Figure S4-.**
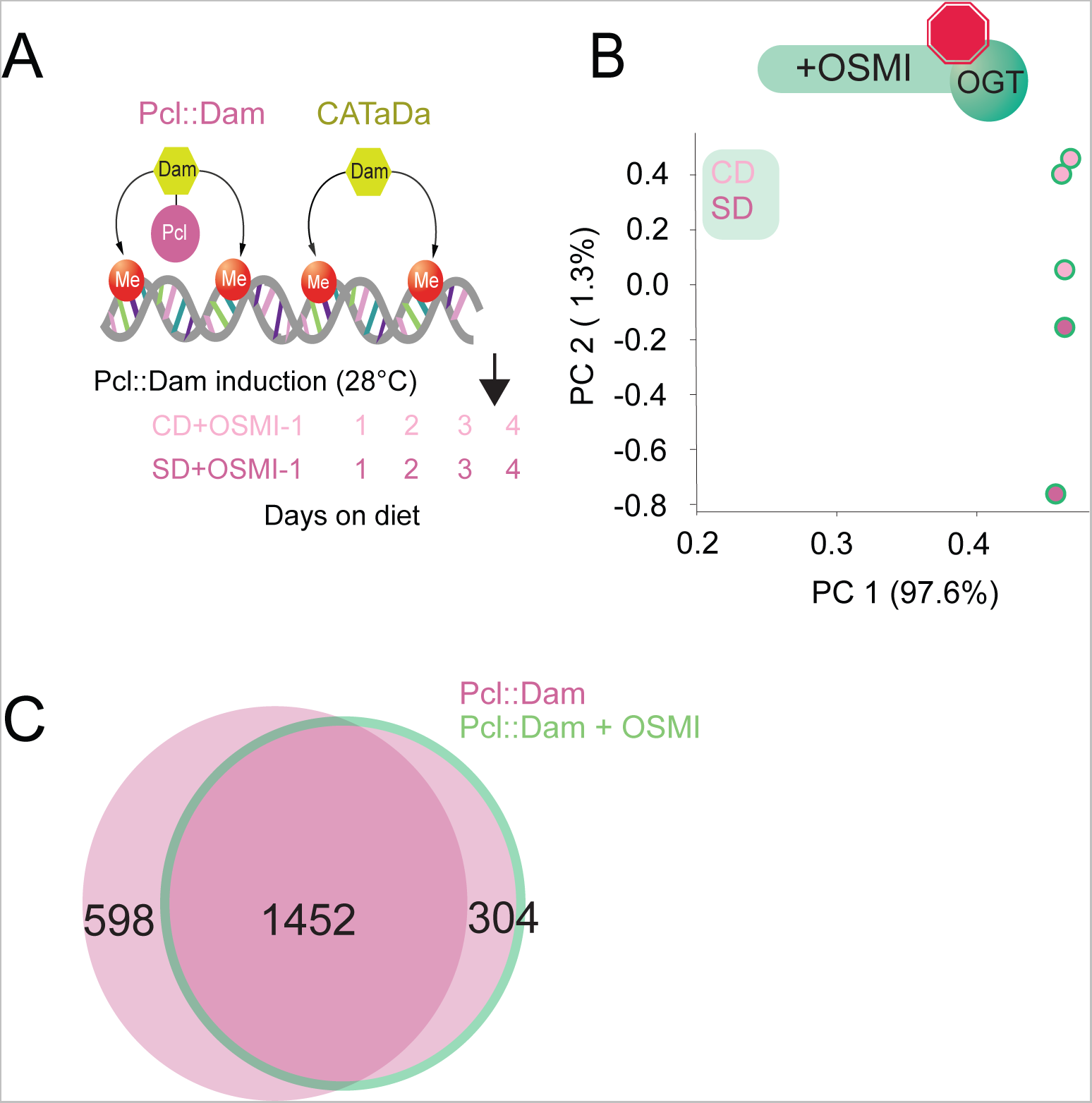
Related to Figure 3, Pcl occupancy at PRE and Pcl peaks with inhibition of OGT activity. (A) Design of the TaDa and CaTaDa experiment in flies fed a CD+OSMI and SD + OSMI. (B) Principal component analysis of normalized log2(*Pcl::Dam/Dam*) flies on CD+OSMI (light pink) or SD+OSMI (dark pink). (C) Overlap of log2(*Dam::Pcl/Dam*) chromatin occupancy peaks in flies with (green outline) or without (pink) OSMI (find_peaks, *q<0.01*).

**Figure S5-.**
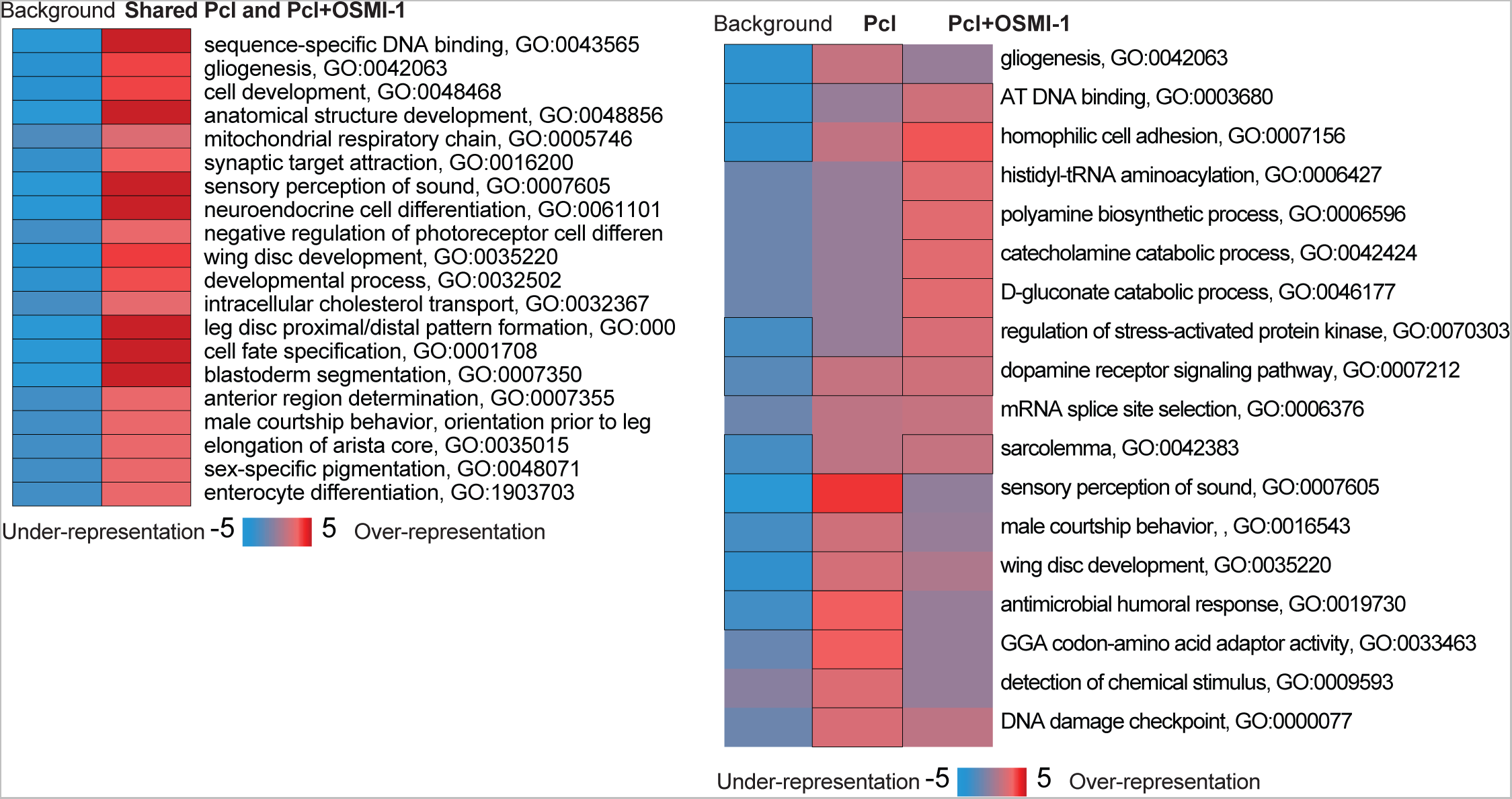
Related to Figure 3, Pathway enrichment analysis of the effects of OSMI on *Dam::Pcl* peaks. iPAGE identification of pathways depleted (blue) or enriched (red) compared to background gene list from the *Dam::Pcl* peaks shared (left) or different (right) between SD/CD with or without OSMI. Scale represents over-representation (red) or under-representation (blue) of genes within a specific bin for the corresponding GO term. Black outlined boxes represent *q<0.05*.

**Figure S6-.**
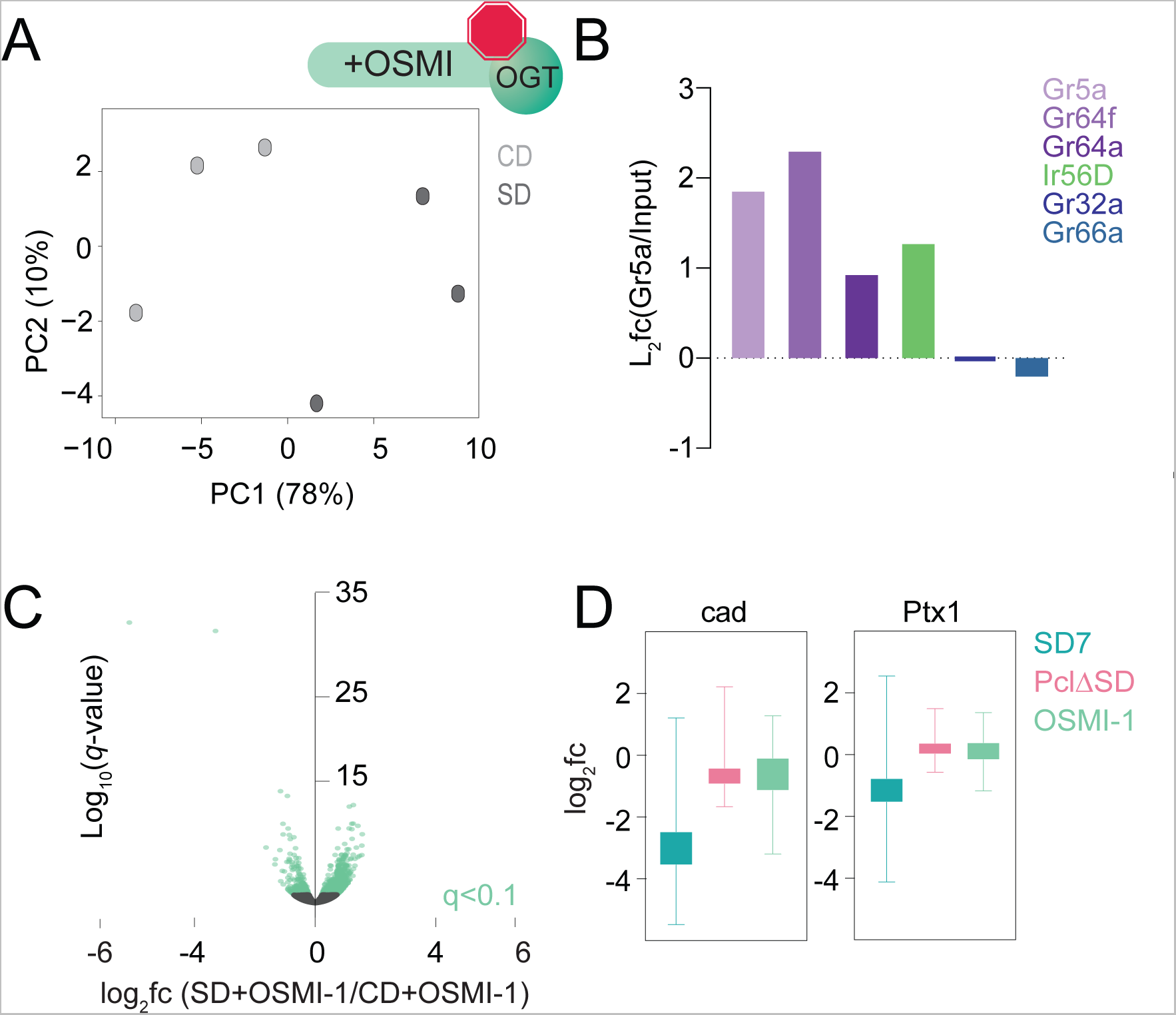
Related to Figure 3, Transcriptional responses to the dietary environment when OGT activity was inhibited. (A) Principal component analysis of *Gr5a > UAS-Rpl3-3XFLAG* flies on a control (light gray) or sugar (dark gray) diet for 7 days supplemented with OSMI-1. (B) Log_2_fold change (l2fc) (Gr5a IP/Input) for *Gr5a, Gr64f, Gr64a, Ir56D, Gr32a, and Gr66a* genes. (C) Volcano plot representing differential expression in the *Gr5a+* neurons of age-matched male *Gr5a>UAS-Rpl3-3XFLAG* flies on a control or sugar diet for 7 days supplemented with OSMI-1. *n*=3 replicates per condition. Genes with *q*<0.1 (Wald test) are in green. (D) Log_2_fold change (l2fc) for candidate gene targets of *Cad* and *Ptx1* at SD7 (teal), *Pcl^c429^ (pink),* and OSMI-1 fed flies at SD7 (green).

**Figure S7-.**
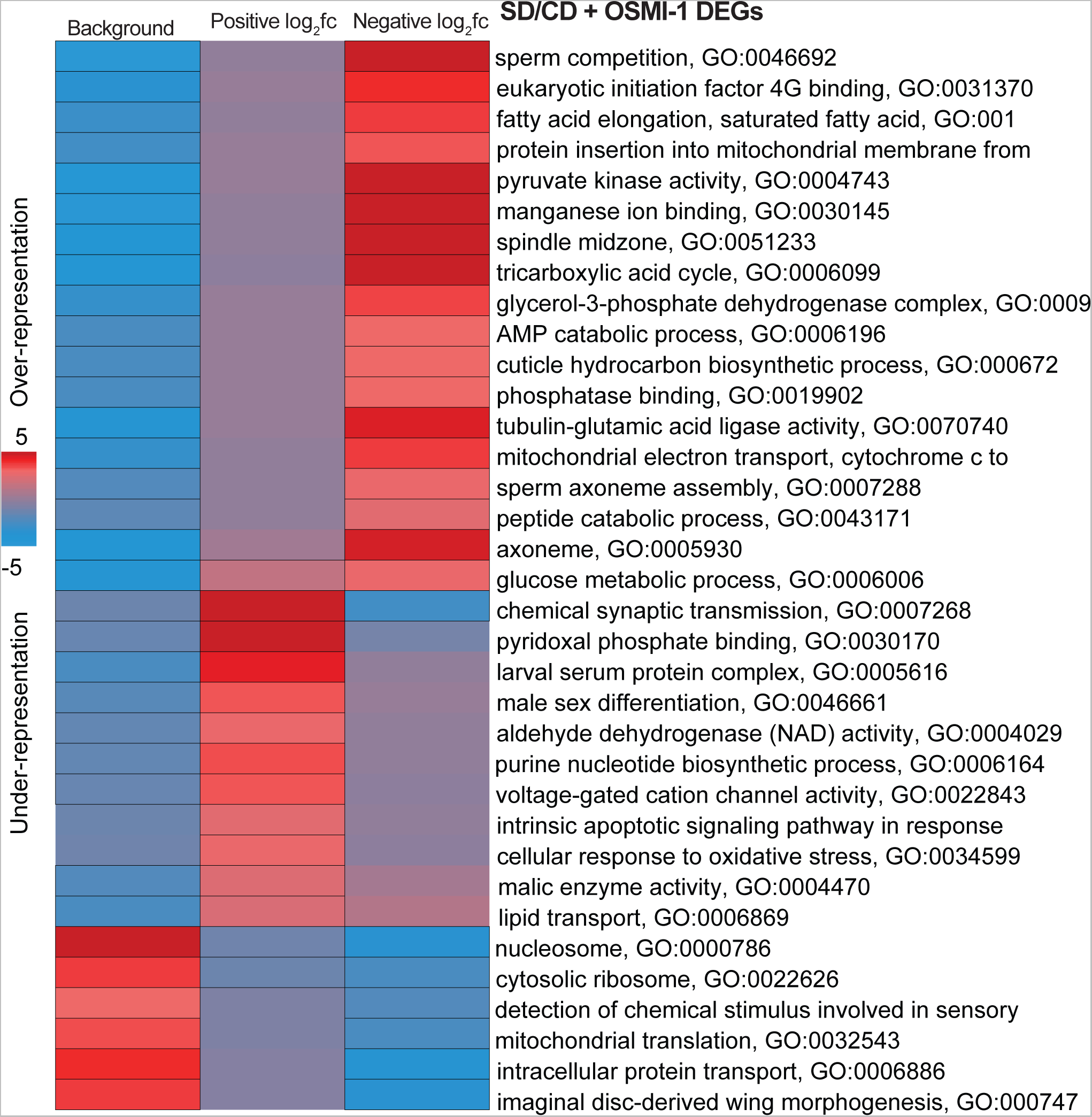
Related to Figure 3, Pathway enrichment analysis of genes reverted or unchanged by OSMI. iPAGE identification of pathways depleted (blue) or enriched (red) compared to background gene list from genes with positive or negative log2 fold changes on SD+OSMI-1. Scale represents over-representation (red) or under-representation (blue) of genes within a specific bin for the corresponding GO term. Black outlined boxes represent q<0.05.

**Figure S8:**
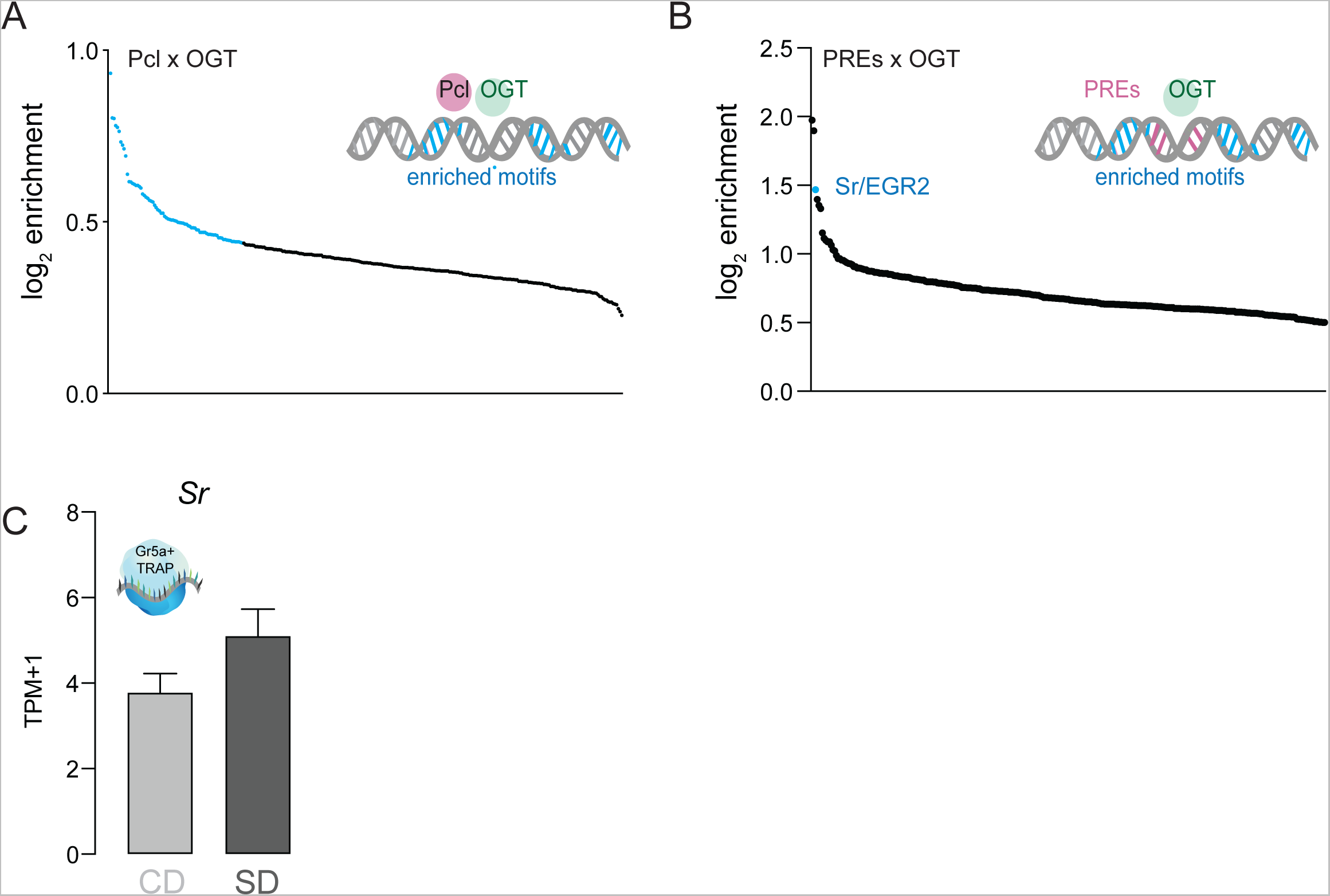
Related to Figure 4, Enrichment analysis of cis-regulatory sites present in OGT and PREs. (A) Log_2_fold enrichment of TF cis-regulatory sites found at OGT + Pcl; blue *q<0.01*. (B) Log_2_fold enrichment of TF cis-regulatory sites found at OGT + PREs. (C) *Sr* mRNA reads normalized as transcripts per million (TPMs) in the Gr5a+ neurons of flies on a CD (light gray) or SD (dark gray) for 7 days, *q>0.01*.

**Figure S9:**
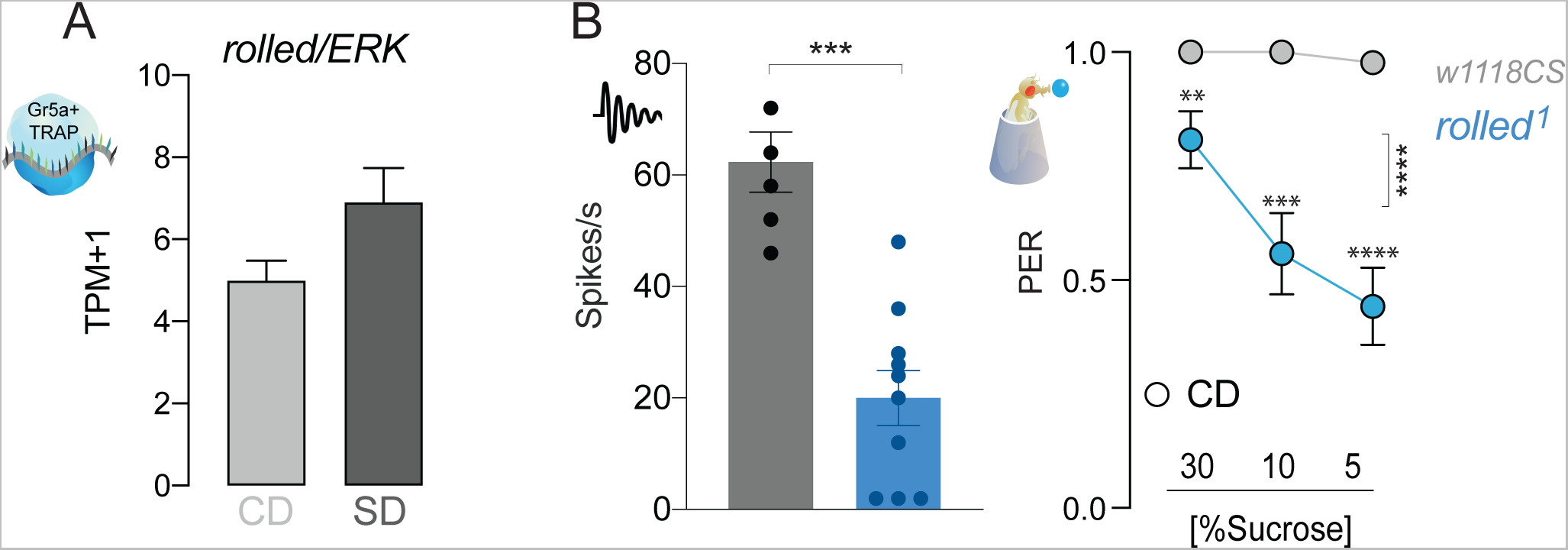
Related to Figure 5, *rl*/ERK is required for taste responses to sugar. (A) *rl* RNA reads normalized as transcripts per million (TPMs) in the Gr5a+ neurons of flies on a CD (light gray) or SD (dark gray) for 7 days. *q>0.01*. (B) (*left*) Averaged neuronal responses to 25 mM sucrose from L-type sensilla in *rl* mutant (blue) and control (gray) flies. n=6-10. Mann-Whitney test: *p*=0.0005. (*right*) Taste responses (*y-axis*) to stimulation of the labellum with 30, 10, and 5% sucrose (*x-axis*) in *rl* mutant and control flies fed a CD. n=26. Two way repeated measure ANOVA, main effect of genotype *p<0.0001*; Dunn’s post-test *\*\*p < 0.01 and ***p<0.001* vs controls.

**Figure S10.**
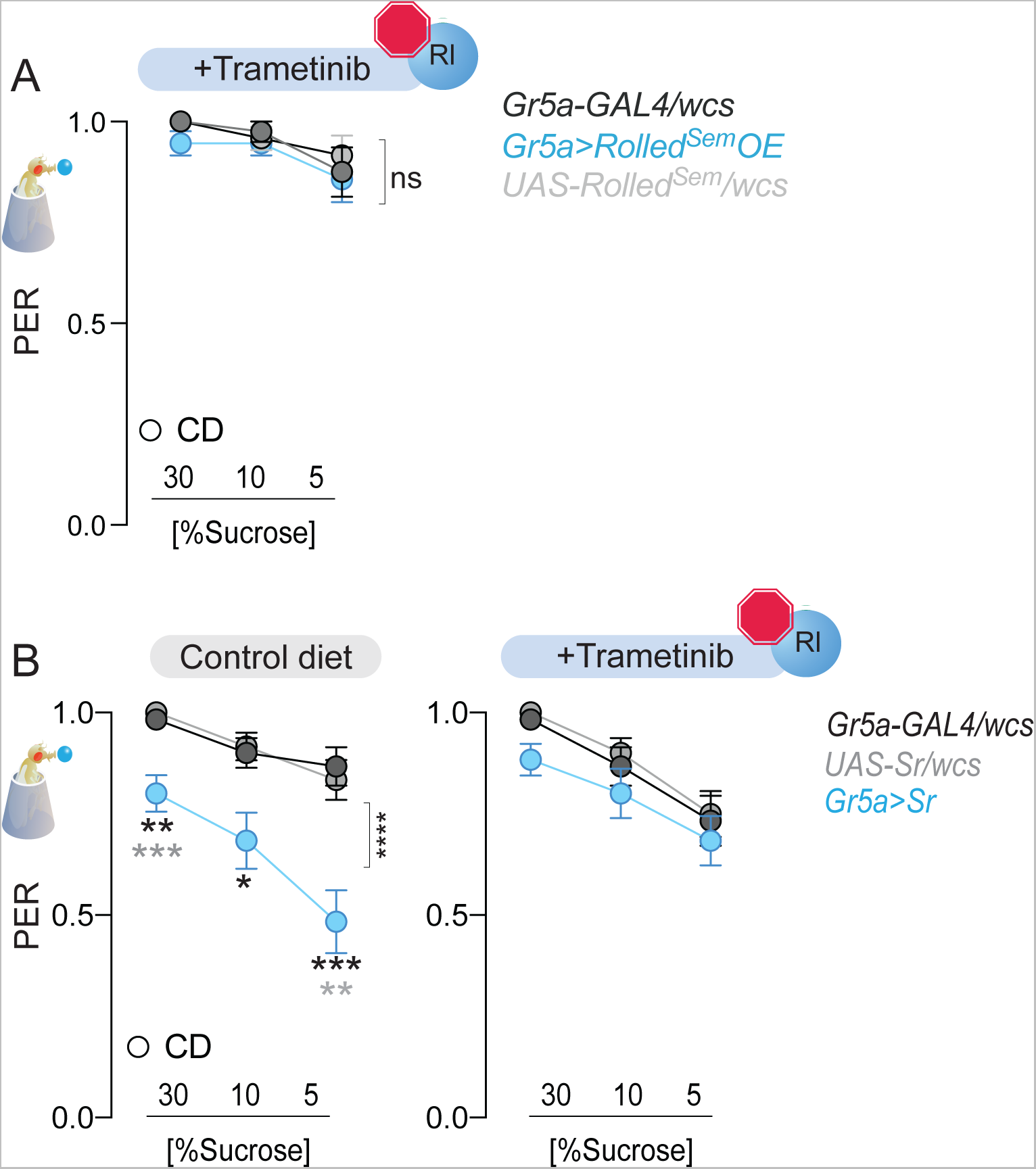
related to Figure 6: Effects of ERK inhibitor Trametinib on genetic manipulations of *Sr* and *rl* levels. (A) Taste responses (*y-axis*) to stimulation of the labellum with 30, 10, and 5% sucrose (*x-axis*) in flies fed a CD+Trametinib (blue) and with overexpression of *rl* (blue or controls in gray) n=24-28. Two-way repeated measure ANOVA, main effect of genotype *p=0.0206* and genotype x concentration *p=0.3671*. (B)Taste responses (*y-axis*) to stimulation of the labellum with 30, 10, and 5% sucrose (*x-axis*) in flies fed a CD with or without Trametinib (blue) and with overexpression of *Sr* (blue, or controls in gray) n=22-38. CD: Two-way repeated measure ANOVA main effect of genotype *p<0.0001* and genotype x concentration *p=0.0037* with Sidak multiple comparison tests, 3*0% p=0.0012, 10% p=0.0002, 5% p<0.0001*; Trametinib: Two way repeated measure ANOVA main effect of genotype *p=0.2192*.

## Notes

### Competing Interest Statement

The authors have declared no competing interest.

### Summary of Updates

This is a major revision; we have added several new data and analyses to better focus the manuscript.

